# FragDockRL: A Reinforcement Learning Method for Fragment-Based Ligand Design via Building Block Assembly and Tethered Docking

**DOI:** 10.1101/2025.08.12.670002

**Authors:** Seung Hwan Hong, Hyunsoo Kim, Se Jin Kim, Soosung Kang

**Affiliations:** Novelism, Seoul 06730, Republic of Korea; College of Pharmacy and Graduate Program in Innovative Biomaterials Convergence, Ewha Womans University, Seoul 03760, Republic of Korea

## Abstract

Efficient exploration of combinatorial chemical space under synthetic constraints remains a central challenge in computational molecular design.

Here, we present FragDock, a molecular design framework that combines building block (BB)-based virtual synthesis with tethered docking guided by a predefined core structure. FragDock defines a structured search space by assembling molecules from synthetically accessible BBs through known chemical reactions and evaluating candidates using tethered docking with a restrained core binding pose. Within this framework, we introduce FragDockRL, a reinforcement learning-based search method that uses docking-score-based rewards and a modified Deep Q-Network (DQN) to guide stepwise molecular growth.

We evaluated FragDockRL on three protein targets, CSF1R, FA10, and VEGFR2, using training-cycle analysis and benchmark comparisons with One-Step Reaction, Random Search, Beam Search, and Monte Carlo Tree Search. FragDockRL progressively enriched molecules with favorable docking scores during learning and generated more cutoff-passing unique molecules than Random Search across all three targets, supporting the benefit of learning-guided prioritization. However, the best-performing search strategy was target-dependent: One-Step Reaction, FragDockRL, and Beam Search each showed advantages in different cases.

Representative molecular case studies showed that selected compounds retained reference-like binding poses while introducing structural variation in peripheral regions. The reaction schemes used commercially available BBs and well-established medicinal chemistry transformations, supporting the synthetic plausibility of the selected compounds. Overall, FragDock provides a flexible framework for synthetically constrained, structure-guided molecular exploration, and FragDockRL offers a learning-guided search mode for productive candidate prioritization under limited generation budgets.

## Introduction

Drug discovery is often perceived as identifying compounds with biological activity. In practice, however, pharmacokinetic properties, safety, and potential side effects are also important considerations for viable drug candidates.^1^ These requirements often impose competing constraints, making it difficult to simultaneously optimize multiple properties in a single molecule. Therefore, increasing the pool of biologically active compounds at an early stage is essential to improve the probability of identifying viable drug candidates. High-throughput screening (HTS) is a widely used experimental approach for the large-scale identification of biologically active compounds and has played a central role in early-stage drug discovery. ^2,3^ In practice, however, the number of compounds that can be experimentally tested is typically limited to the order of hundreds of thousands.

By contrast, computer-based virtual screening enables the screening of chemical libraries containing hundreds of millions to billions of molecules.^4^ In such large-scale virtual screening, molecular docking is widely used to predict ligand binding poses and estimate binding affinity as a proxy for biological activity. ^5–8^ Nevertheless, evaluating hundreds of millions of compounds still requires substantial computational resources.

To reduce the computational burden, various heuristic search strategies have been explored to efficiently navigate large chemical spaces, including greedy search^9^ and Thompson sampling.^10^ In addition, machine learning (ML) approaches have also been developed to accelerate virtual screening, for example by training ligand-based ML models on a docked subset of a large library to predict docking scores for the remaining molecules^11,12^ or by guiding molecular generation using docking-derived rewards. ^13,14^

Search strategies are not limited to computational algorithms but can also incorporate principles from medicinal chemistry. For instance, V-SYNTHES^15^ reduces the search space through synthon-based hierarchical enumeration, reflecting the medicinal chemistry strategy of exploring substituent combinations around reaction scaffolds. By evaluating and expanding selected scaffold–synthon combinations, the search space can be effectively reduced. Despite these advances, efficiently exploring the combinatorial space of synthetically accessible building blocks (BBs) under structural constraints imposed by protein binding sites remains a major challenge.

Understanding the relationship between the structural features of ligand binding sites in proteins and those of biologically active molecules can help reduce the effective search space. For example, kinase inhibitors often contain a core structure known as a hinge binder that interacts with the hinge region and plays a key role in binding affinity. Many compounds targeting the same protein share similar core structures, a feature exploited in fragment-based drug design (FBDD).^16^ In FBDD, a core structure is fixed while surrounding substituents are optimized to improve binding. If a substituent does not fit the binding pocket, the molecule cannot bind effectively, resulting in loss of activity. This strategy can also be translated into computational virtual screening. Tethered docking is a docking approach in which a subset of atomic coordinates is restrained while allowing the remaining parts of the molecule to be optimized.^17^ Such approaches are commonly used to guide the search toward ligand binding modes consistent with a reference core binding mode.

In addition to structural considerations, chemical synthesis significantly constrains the exploration of chemical space. The synthesis of complex molecules often requires multi-step reactions, and not all theoretically generated compounds are synthetically accessible. In early-stage hit identification, medicinal chemists typically design molecules from commercially available BBs using well-established chemical reactions. This BB—based paradigm has also been adopted in computational approaches. For example, large virtual libraries such as the Enamine REAL database^18^ are constructed by enumerating combinations of synthetically accessible BBs according to known reaction rules, enabling the exploration of billions of compounds while maintaining a high degree of synthetic feasibility.

Building on these concepts, we introduce FragDock, a molecular search framework that combines BB-–based molecular assembly with tethered docking guided by a predefined core structure. FragDock restricts the search space to synthetically accessible molecules by assembling BBs through known chemical reactions, effectively generating candidates via virtual synthesis. In parallel, the core binding pose is restrained, and candidate molecules are evaluated using tethered docking based on its three-dimensional reference structure. Within this framework, candidate molecules are constructed and evaluated, while their exploration is governed by a search strategy.

In this study, we develop FragDockRL, a search strategy within the FragDock framework based on a modified Deep Q-Network (DQN), a reinforcement learning (RL) algorithm.^19^ At each step, FragDockRL operates on a partially constructed molecule and selects the next BB and reaction to extend it, framing molecular assembly as a sequential decision process. Compared to conventional search methods that rely on explicit evaluation of generated candidates, learning-based approaches can learn structural features of BBs and estimate action values for unexplored candidates, thereby potentially enabling more efficient exploration of the search space. An overview of the proposed FragDockRL framework is shown in Figure 1.

**Figure 1:**
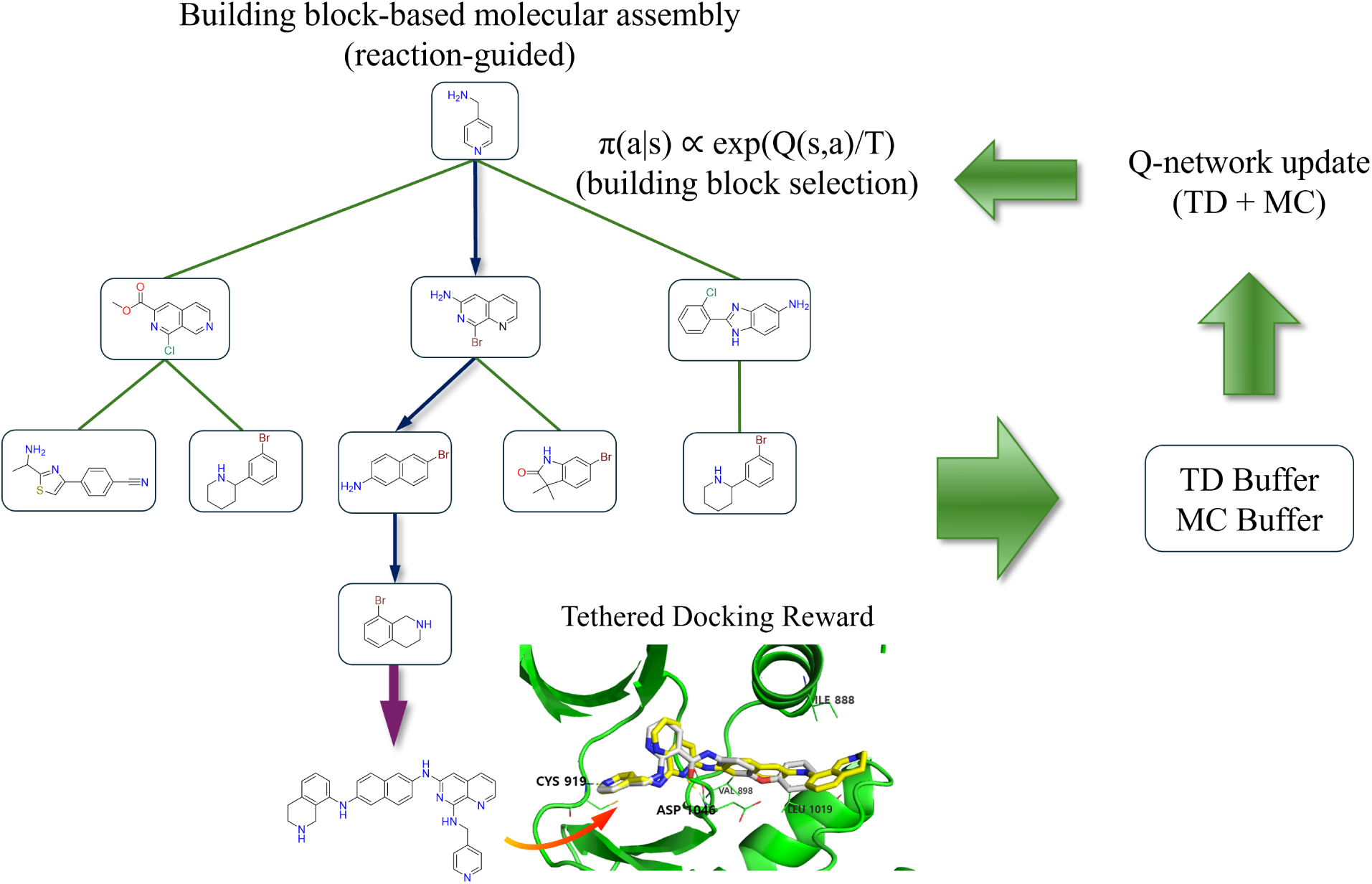
Overview of the FragDockRL framework. Molecules are generated through reaction-based BB assembly from an initial BB that contains the core structure used for tethered docking restraints. Additional BBs are sequentially selected using Q-values estimated by the Q-network. The generated molecules are evaluated using tethered docking with a restrained core pose, and the resulting docking scores are used to guide the search. Trajectories are stored in temporal-difference (TD) and Monte Carlo (MC) replay buffers, and the Q-network is updated using these buffers.

FragDockRL was evaluated on three target proteins—CSF1R, FA10, and VEGFR2— for structure-guided candidate generation. To systematically assess its search behavior, we compared FragDockRL with four baseline methods implemented within the same Frag-Dock framework: One-Step Reaction, Random Search, Beam Search, and Monte Carlo Tree Search.^20,21^ Together, these experiments assess the utility of FragDockRL for exploring syn-thetically accessible molecular candidates with favorable predicted binding poses under limited generation budgets, without requiring large-scale labeled biological activity datasets.

## Methods

### Building Block (BB) and Reaction Space

The BB library and reaction rules defining the chemical search space in FragDock are as follows. We utilized 58 reaction SMIRKS reported in the study by Hartenfeller et al.^22^

A total of 124,180 BBs provided by Enamine^18^ were collected. Prior to filtering and reaction matching, salt fragments and counterions were removed from the BB SMILES, and the major molecular component was retained as the representative BB structure. BBs with molecular weights exceeding 300 were excluded, resulting in 113,515 selected BBs. Each BB was matched to applicable reaction rules based on its functional groups, defining the set of valid reactions during molecule assembly. For computational efficiency, applicable reaction rules were precomputed for each BB by matching the reactant-side SMARTS patterns of the reaction SMIRKS to the BB structures using RDKit^23^ substructure matching.

### Target Proteins and Structural Data

Three target proteins—CSF1R, FA10, and VEGFR2—were selected to evaluate the molecular generation and docking performance of FragDockRL across structurally distinct binding pockets with well-characterized co-crystallized ligand structures.

CSF1R (Colony Stimulating Factor 1 Receptor): A receptor tyrosine kinase involved in the proliferation and differentiation of macrophages and myeloid cells. It is an important target in the development of therapies for inflammatory diseases and certain cancers.^24^

FA10 (Coagulation Factor Xa): A protease enzyme central to the blood coagulation process, serving as a key target in the development of anticoagulants for the prevention and treatment of thrombosis.^25^

VEGFR2 (Vascular Endothelial Growth Factor Receptor 2): A receptor involved in angiogenesis, playing a key role in tumor blood vessel formation and growth. Inhibition of VEGFR2 is a major strategy in cancer therapy, making it a primary target in the development of anti-cancer drugs.^26^

The target protein structures were selected based on the PDB IDs provided in the DUD-E dataset,^27^ and the corresponding PDB files were downloaded from the Protein Data Bank (PDB).^28^ For each structure, the protein and co-crystallized ligand were separated. The protein structure was prepared using PDBFixer^29^ by repairing missing atoms, assigning protonation states, and adding hydrogen atoms. The co-crystallized ligand was used as the reference ligand for defining the core conformation and the docking region. The corresponding binding pocket structures, co-crystallized ligands, core structures, and initial BBs are shown in Figure 2. The PDB IDs, reference ligand IDs, and core SMILES used for each target are summarized in Table 1.

**Figure 2:**
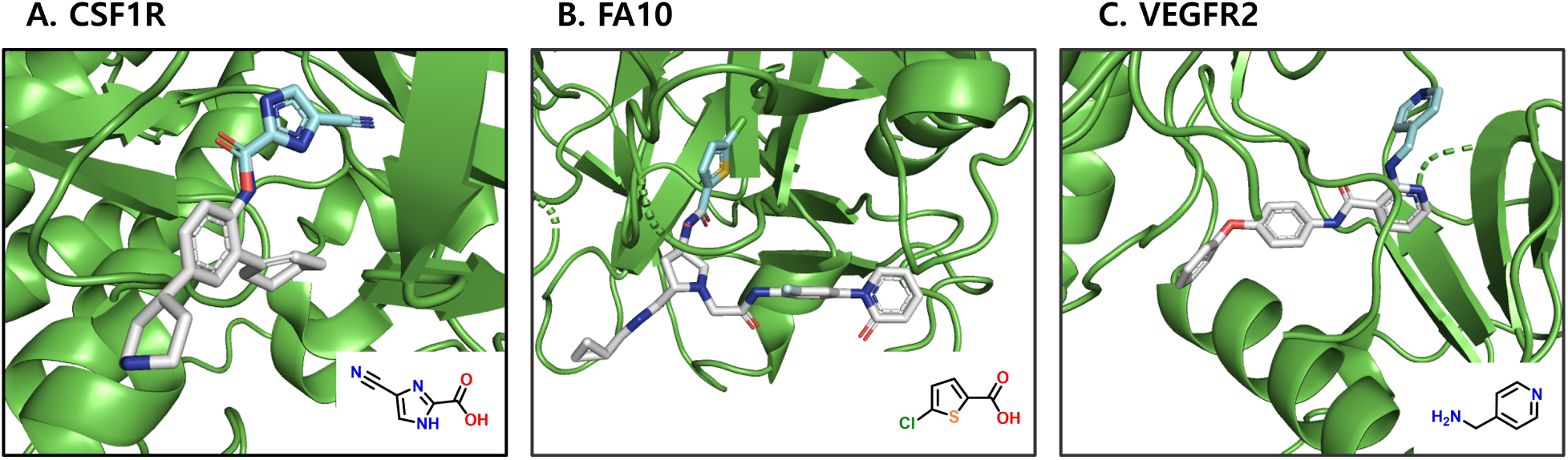
Binding pocket structures and initial BBs for the three targets: (A) CSF1R, (B) FA10, and (C) VEGFR2. Protein crystal structures are shown in green, co-crystallized ligands in light gray, and core structures used for tethered docking restraints in cyan. Insets show the initial BBs used as the initial states in FragDockRL.

**Table 1:** Target protein structures and molecular inputs used for FragDockRL. The co-crystal docking score is reported in kcal/mol and was computed by single-point rescoring of the crystallographic ligand pose without additional conformational sampling. The core SMILES denotes the core structure used for the tethered docking restraint, whereas the initial BB SMILES denotes the initial building block used for molecule generation.

| Parameter | CSF1R | FA10 | VEGFR2 |
| --- | --- | --- | --- |
| PDB ID | 3KRJ | 2VWM | 2P2I |
| Co-crystal ligand ID | KRJ | LZI | 608 |
| Co-crystal docking score | -8.561 | -10.136 | -11.422 |
| Core SMILES | N#Cc1nc(nc1)C | C(=O)c1ccc(Cl)s1 | Cc1ccncc1 |
| Initial BB SMILES | N#Cc1nc(nc1)C(=O)O | OC(=O)c1ccc(Cl)s1 | NCc1ccncc1 |

### Tethered Docking

Prior to docking, an initial conformer was generated for each molecule using RDKit^23^ without imposing constraints on the core. The generated conformer was then aligned to the reference core structure based on substructure matching, followed by a coordinate-based alignment of the conformer. When multiple valid matches were identified, tethered docking was performed for each match independently.

Tethered docking was performed using rDock^8^ with 10 generated conformers per ligand. The binding cavity was defined with a radius of 8.0 Å. Ligand sampling was conducted under tethered constraints, with translational and rotational degrees of freedom restricted (MAX_TRANS = 1.0, MAX_ROT = 30.0), while dihedral angles were allowed to vary freely. Additional parameters were kept at their default values unless otherwise specified.

Each conformer was subsequently energy-minimized using SMINA^7^ with the --minimize option only, which performs local energy optimization without additional conformational sampling. Thus, conformational sampling was primarily handled by rDock, while SMINA was used solely for local refinement. For compatibility with SMINA, both protein and ligand structures were converted from PDB to PDBQT format using AutoDockTools.^30^

The docking score computed using SMINA (in kcal/mol) is referred to as the raw docking score, where more negative values indicate stronger binding. To ensure consistent handling of failed or unsuitable docking cases, the score was post-processed. Specifically, docking failures, conformers with a docked-core root-mean-square deviation (RMSD) greater than 0.5 Å relative to the reference core structure, excessively high unfavorable scores, and molecules excluded from docking based on predefined thresholds were assigned a fixed value of 3 kcal/mol. Molecules with more than 12 rotatable bonds, more than 60 heavy atoms, or a molecular weight greater than 650 were excluded from docking. These criteria were applied as loose filtering criteria rather than strict drug-likeness constraints.

Among all conformers across all valid core matches, the conformer with the lowest post-processed value was selected as the final result for the molecule. In the following sections, this post-processed value is referred to as the docking score.

### FragDock Framework

FragDock is a molecular design framework that generates candidate molecules through BB–based assembly and evaluates them using tethered docking with a predefined core structure. The framework defines a structured chemical search space by combining synthetically accessible BBs under predefined reaction rules.

Molecule construction is performed sequentially via virtual synthesis. The process begins with an initial BB containing the core structure, and additional BBs are iteratively incorporated through compatible reactions. At each step, applicable reaction rules are determined based on the current molecular structure, and only BBs capable of participating in those reactions are considered as valid candidates.

In practice, reaction applicability is determined by matching the reactant-side SMARTS patterns of each reaction template, and product generation is performed by applying the corresponding reaction SMIRKS using RDKit. Reactions that can produce multiple structurally distinct products from the same BB combination are considered invalid and are not allowed in the assembly process. This constraint avoids ambiguity in state transitions by ensuring a well-defined and deterministic mapping between actions and resulting molecular states.

The assembly process continues until a predefined maximum number of steps is reached, no further valid reactions are available, or a special termination BB (index 0) is selected. The resulting molecule is then evaluated using tethered docking, where the binding pose of the core structure is restrained to ensure consistent evaluation across generated molecules.

### FragDockRL

FragDockRL is a reinforcement learning-–based search strategy within the FragDock framework, in which a Q-network evaluates candidate BB additions during sequential molecular assembly. At each step, an action corresponding to a BB addition is selected to extend the current molecule, and the terminal molecule generated at the end of an episode is evaluated using tethered docking to provide the final reward signal. Through repeated interactions, the Q-network learns to estimate action values, which are used to guide the selection of promising molecular constructions.

#### Formulation

In FragDockRL, an episode is defined as a sequence of state–action transitions that begins with the initial BB containing the core structure and ends at a terminal molecule. The state is defined as the current molecular structure, and the action corresponds to selecting a BB to be attached or a termination action. The assembly process proceeds sequentially until a termination action is selected or no further valid reactions are available. The transition is defined by applying the selected BB through a reaction. If the reaction yields a single valid product, the state is updated to the resulting molecule. Otherwise, the transition is treated as invalid; the state remains unchanged, and a penalty is applied.

The number of BB additions is limited to a predefined maximum (set to 3 in this study). Once this limit is reached, only the termination action is allowed. In our implementation, the termination action is represented by a dedicated BB index (index 0).

The reward consists of step-wise penalties incurred during an episode and a final reward assigned at the terminal step. At each step within an episode, a penalty of −2 is assigned if the selected action results in an invalid reaction, where the state remains unchanged. Valid transitions receive no step-wise reward. At the end of an episode, the final reward is defined based on the post-processed docking score described in the Tethered Docking subsection. The docking score, where more negative values indicate stronger predicted binding, is converted into a dimensionless reward by negating its value. Thus, higher reward values correspond to stronger predicted binding affinity. The total return is defined as the sum of all step-wise penalties and the final reward.

#### Search (Molecule Generation)

Based on the formulation above, molecular construction proceeds as follows. At each step within an episode, an action is selected according to the policy and applied via the corresponding reaction. The process continues until a termination action is selected or no further valid actions are available, resulting in a complete molecule.

We adopt a Q-value-–based formulation, as the feasible action set varies with the current state. The Q-function naturally evaluates only the valid candidates at each step. To select an action given the current state, a policy is required. In FragDockRL, the policy is derived from an action-value function *Q*(*s, a*), which estimates the value of selecting action *a* in state *s*. The Q-function is approximated by a neural network, which is described in detail in the Q-network and Training subsection. To reduce the size of the action space and improve computational efficiency, candidate BBs are first filtered based on the valid reaction rules, and Q-values are evaluated only over the resulting feasible action set A(*s*).

The stochastic policy is derived from the Q-function via a Boltzmann distribution, as shown in Eq. 1:

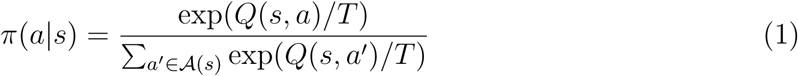

The temperature parameter *T* is gradually decreased during training to balance exploration and exploitation. Specifically, at cycle *k* (each consisting of a batch of molecule generation followed by training), the temperature is defined by Eq. 2:

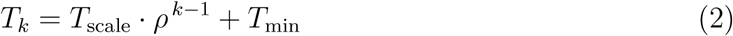

where *T*_scale_ controls the decaying component of the temperature, *ρ* is the temperature decay factor, and *T*_min_ is the minimum temperature. In this study, *T*_scale_ = 0.80, *ρ* = 0.99, and *T*_min_ = 0.20, giving an initial temperature of *T*_scale_ + *T*_min_ = 1.00. The minimum temperature term *T*_min_ prevents overly deterministic action selection at low temperatures and helps maintain diversity. The temperature parameters were selected based on preliminary trial runs.

An action is selected by sampling from the policy *π*(*a*|*s*). This stochastic selection enables exploration of diverse molecular structures while favoring actions with higher Q-values.

#### Data Generation and Replay Buffer Construction

Unlike standard DQN settings, which often involve longer trajectories and provide abundant training signals per episode, FragDockRL generates relatively short episodes with sparse and delayed rewards, where meaningful feedback is primarily obtained at the terminal step. In addition, reward evaluation based on docking is computationally expensive, making step-wise updates inefficient. Therefore, FragDockRL organizes the training process into cycles, in which multiple episodes are generated, evaluated in batch, and then used to update the Q-network. This formulation also enables efficient parallelization of docking and improves computational efficiency. In addition, to handle the sparse and terminal-dominated reward structure, FragDockRL incorporates both temporal-difference (TD) and Monte Carlo (MC) learning signals.

At each cycle, a batch of molecular construction episodes is first generated using the current policy. Each episode corresponds to the construction of a single molecule, and episodes are generated sequentially until the batch is complete. In this study, 200 molecules are generated per cycle. After generation, duplicate molecules are merged based on their SMILES representations before final reward evaluation. Docking is then performed in parallel for the resulting set of unique molecules, and the corresponding final rewards are computed. The replay buffers are subsequently updated using the collected transition-level data and trajectory-level returns.

Specifically, the transition-level data and trajectory-level returns are stored separately in two replay buffers. The TD buffer stores transition-level data (*s_t_, a_t_, r_t_*_+1_*, s_t_*_+1_) and is used for bootstrapped updates, while the MC buffer stores trajectory-level data (*s_t_, a_t_, g_t_*), where *g_t_* is the discounted return defined in Eq. 3.

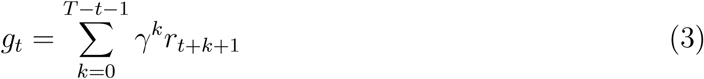

The MC buffer stores trajectory-level return targets that include the final reward.

The TD buffer accumulates transition data across cycles with a maximum capacity of 40,000 samples. In contrast, the MC buffer is constructed within each cycle using the trajectories generated in that cycle and is not carried over across cycles. This cycle-local construction is used to compute MC returns from trajectories generated by the current policy.

A total of 100 cycles are performed, each consisting of molecule generation and subsequent Q-network training using the resulting rewards, corresponding to up to 20,000 generated molecules.

#### Q-network and Training

The Q-network estimates the action-value *Q*(*s_t_, a_t_*) given a state–action pair. Both the current molecule (state) and the candidate BB (action) are represented as 1024-bit Morgan fingerprints^31^ with radius 2. These inputs are processed by a shared encoder consisting of a multi-layer perceptron with three hidden layers, producing 512-dimensional latent representations. The encoder is shared between the state and action inputs. The latent representations of the state and action are concatenated and fed into a second multi-layer perceptron, which outputs a scalar value corresponding to *Q*(*s_t_, a_t_*), as illustrated in Figure 3. All hidden layers use ReLU activation functions. A separate target network *Q*_target_ is maintained alongside the online Q-network *Q*, and is used for computing TD targets.

**Figure 3:**
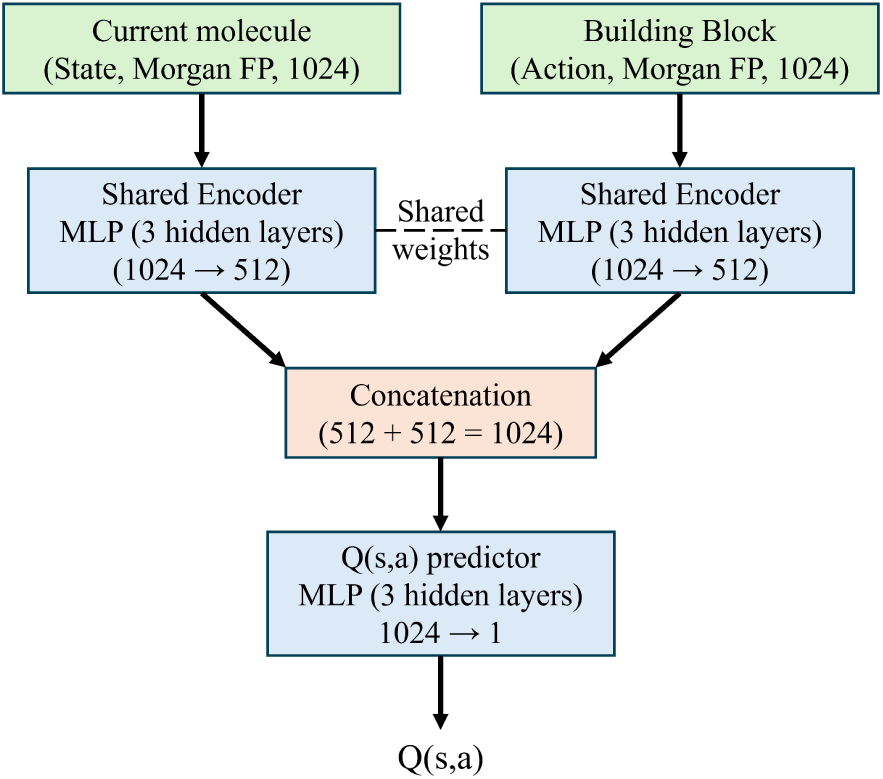
Q-network architecture in FragDockRL. State and action representations are encoded by a shared network, concatenated, and used to estimate the action-value *Q*(*s, a*).

Following data generation, the Q-network is trained at each cycle using samples from the replay buffers. The Q-network is updated for 12 iterations per cycle using mini-batches of size 128. During training, each mini-batch is constructed by sampling from both the TD and MC buffers, with the numbers of TD and MC samples determined adaptively.

The number of MC samples used within a training phase is capped at 50% of the total training samples. When the MC buffer contains enough samples to reach this upper bound, each mini-batch contains an equal number of TD and MC samples. Otherwise, all available MC samples are distributed across iterations, resulting in fewer MC samples per mini-batch and a correspondingly larger number of TD samples. TD samples are randomly drawn from the TD buffer at each iteration, while MC samples are shuffled once and then consumed sequentially across iterations.

Based on the sampled mini-batches, the Q-network is trained using the loss terms defined below. The TD target is given by Eq. 4:

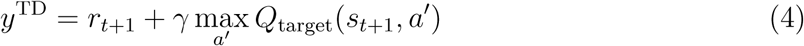

where *γ* = 0.9 is the discount factor. The TD loss is defined in Eq. 5:

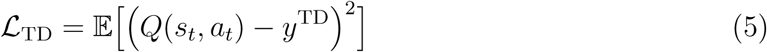

The MC loss is defined in Eq. 6:

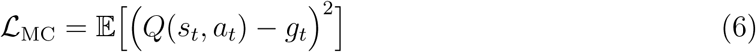

where *g_t_* is the discounted return defined in Eq. 3. The final loss is computed as a weighted average of the TD and MC losses based on the number of samples in each mini-batch, as shown in Eq. 7:

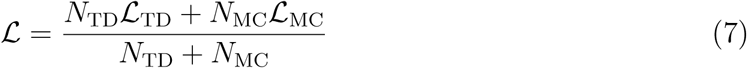

The network parameters are optimized using the Adam optimizer with a learning rate of 0.001.^32^ The weights are initialized using Xavier normal initialization,^33^ and the biases are initialized from a normal distribution with zero mean and unit variance. A target network is maintained and updated using a soft update scheme with a smoothing coefficient *τ* = 0.005 after each optimization step.

We consider both TD and MC updates for training the Q-network, and also evaluate a TD-only variant by disabling MC updates.

### Baseline Methods

To evaluate the effectiveness of FragDockRL, we consider several baseline methods, including three search-based approaches—–Random Search, Beam Search (BS), and Monte Carlo Tree Search (MCTS)—–as well as a One-Step Reaction baseline. Since these methods differ in their exploration and selection strategies, a consistent evaluation criterion is required. In this work, we use the number of docking evaluations as the primary metric, as docking constitutes the dominant computational cost.

The search-based methods are compared under a docking budget of up to 20,000 evaluations, consistent with the maximum budget used in FragDockRL. For BS, this corresponds to 19,998 docking evaluations due to its step-wise allocation. In contrast, the One-Step Reaction baseline exhaustively evaluates all molecules that can be generated by a single reaction from the core structure.

All baseline methods are implemented within the same FragDock framework, and all settings are identical to those of FragDockRL unless otherwise specified.

#### One-Step Reaction

This baseline performs exhaustive enumeration of all molecules that can be generated by a single reaction from the core structure. This setting corresponds to a single-step synthesis scenario and provides a reference for comparing exhaustive single-step evaluation with multi-step search strategies.

#### Random Search

Random Search follows the same environment and reaction rules as FragDockRL but selects actions uniformly at random without using a Q-network or any learning. Molecules are generated step-by-step until the predefined budget is reached.

#### Beam Search (BS)

We employ a stochastic BS to reduce the computational cost of evaluating all possible actions. At each step, a fixed number of candidate actions (*N*_expand_ = 6666) are randomly sampled and applied to the current molecule to generate new molecules, which are then evaluated via docking. The top-*k* molecules (*k* = 10) are retained for further expansion. This procedure is repeated for three steps, resulting in a total of 19,998 docking evaluations. This approach approximates standard BS under a limited docking budget.

#### Monte Carlo Tree Search (MCTS)

We apply a standard MCTS algorithm within the FragDock framework to explore the molecular assembly space. The root node corresponds to the core molecule, and each node represents a molecular state, with edges corresponding to reaction-based BB additions.

At each iteration, nodes are selected using an upper confidence bound (UCB) criterion, followed by expansion and rollout using a random policy. Each rollout proceeds up to the maximum number of steps, and the resulting molecule is evaluated via docking as the terminal reward. The reward is then backpropagated to update the node statistics.

## Results and discussion

FragDockRL was evaluated on three targets: CSF1R, FA10, and VEGFR2, following the experimental protocol described in Methods. Its performance was compared with baseline methods implemented within the same FragDock framework. Each method was independently repeated five times for each target, except for the One-Step Reaction baseline, which was performed once because it is deterministic and does not involve repeated stochastic sampling. We first analyze the learning behavior of FragDockRL across cycles, and then present benchmark comparisons and representative molecular case studies.

### Learning Behavior Across Cycles

Before analyzing the cycle-dependent results, we note that FragDockRL is affected by both Q-network training and the temperature-controlled exploration schedule. Because the objective is to identify molecules with favorable docking scores within a limited docking budget, rather than to fully converge the Q-function, we examined whether the selected schedule improved docking scores while maintaining molecular diversity. We separately trained and compared two FragDockRL variants: TD-only, which used only the TD loss, and TD+MC, which used both the TD and MC losses. For each variant, we monitored two cycle-dependent metrics: the average docking score of generated molecules and the cumulative numbers of unique molecules and unique Bemis–Murcko scaffolds^34^ for each target (Figure 4).

**Figure 4:**
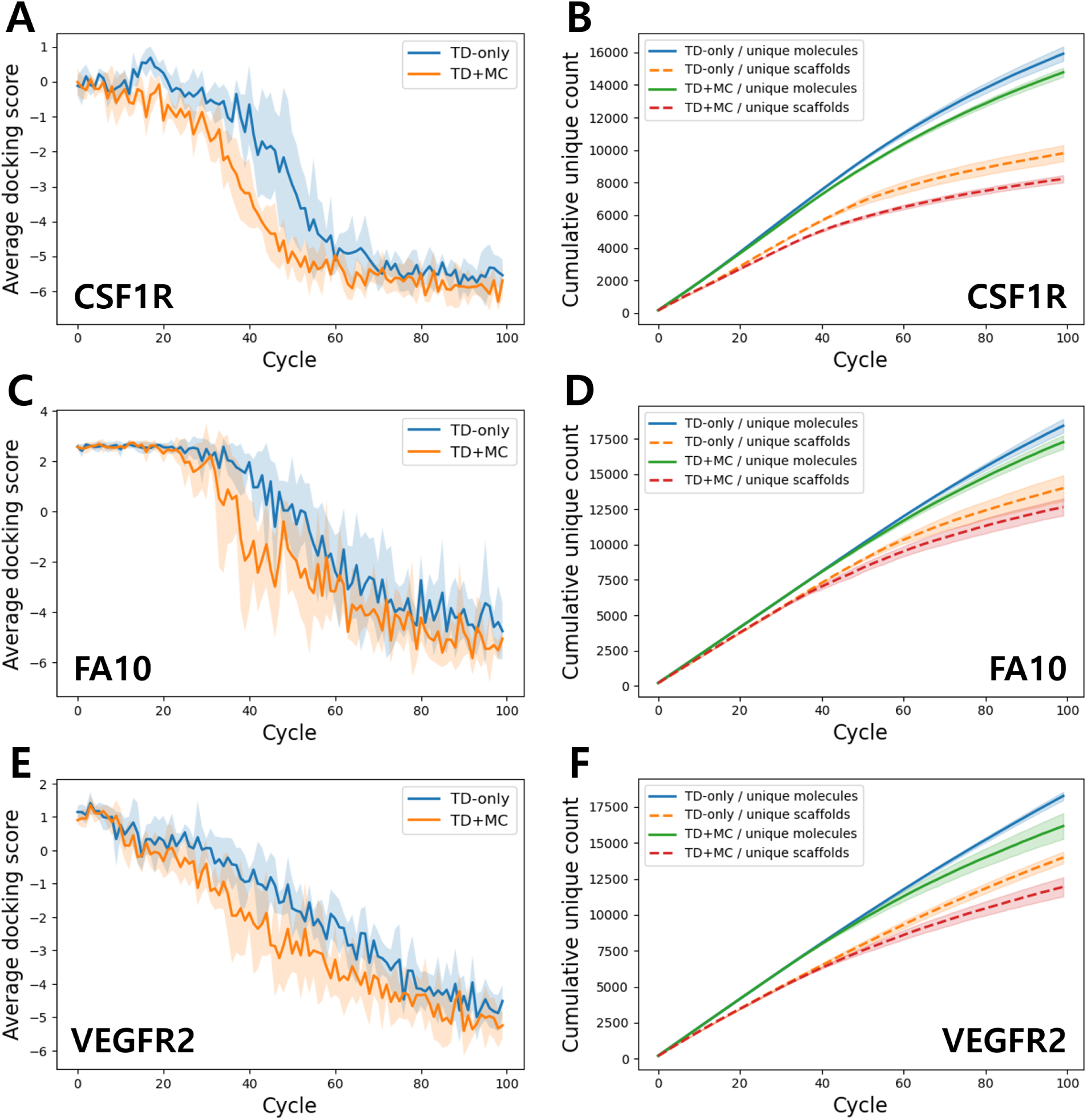
Cycle-dependent learning behavior of FragDockRL. Panels A, C, and E show the cycle-wise mean docking score, reported in kcal/mol, of generated molecules for CSF1R, FA10, and VEGFR2, respectively. Panels B, D, and F show the cumulative numbers of unique molecules and unique Bemis–Murcko scaffolds discovered across cycles for CSF1R, FA10, and VEGFR2, respectively. Solid lines represent the mean of five independent runs, and shaded regions indicate the standard deviation.

For all three targets, the average docking score decreased as the learning cycles progressed (Figure 4A,C,E). This indicates that Q-network training progressively biased action selection toward BB additions that generated molecules with more favorable docking scores. The decrease was generally faster for TD+MC than for TD-only, suggesting that the inclusion of the MC loss accelerated this shift in Q-guided molecule generation.

The detailed profiles of docking-score improvement differed among the targets. For CSF1R, the average docking score decreased only modestly during the early cycles, followed by a sharper decrease in the middle of the run, after which the improvement became more gradual. For FA10, the average docking score remained high, above 2 kcal/mol, during the early cycles and then started to decrease more clearly in subsequent cycles. This high initial average score indicates that docking failures, which were assigned a score of 3 kcal/-mol, were dominant in the early cycles. Compared with CSF1R and FA10, VEGFR2 showed a more gradual and nearly continuous decrease throughout the 100 cycles. These different improvement profiles may reflect target-dependent differences in the docking landscape and the accessible molecular space around each initial state.

The cumulative numbers of unique molecules and unique scaffolds increased throughout the cycles for all targets, but the rate of increase gradually decreased in later cycles (Figure 4B,D,F). This trend can be partly attributed to the decreasing temperature schedule, which reduces random exploration over cycles and increases the influence of Q-guided action selection. Compared with TD-only, TD+MC generally showed slower growth in the numbers of unique molecules and unique scaffolds. Because the same temperature schedule was used for both variants, this result suggests that the extent of Q-network training also contributed to the narrowing of the explored chemical space.

Although the average docking score decreased over learning cycles, the objective of Frag-DockRL is not only to improve the overall docking score distribution but also to discover diverse molecules with docking scores lower than a predefined cutoff. We therefore applied target-specific docking score cutoffs and selected only molecules with docking scores lower than the cutoff values. The cutoff values were set to −8, −9, and −11 kcal/mol for CSF1R, FA10, and VEGFR2, respectively, based on the redocking scores of the co-crystallized ligands (Table 1).

Figure 5 shows the cycle-dependent discovery of molecules and scaffolds passing these docking score cutoffs. Panels A, C, and E show the cumulative numbers of unique molecules and unique scaffolds for CSF1R, FA10, and VEGFR2, respectively. Panels B, D, and F show the number of unique scaffolds discovered in each cycle for the corresponding targets.

**Figure 5:**
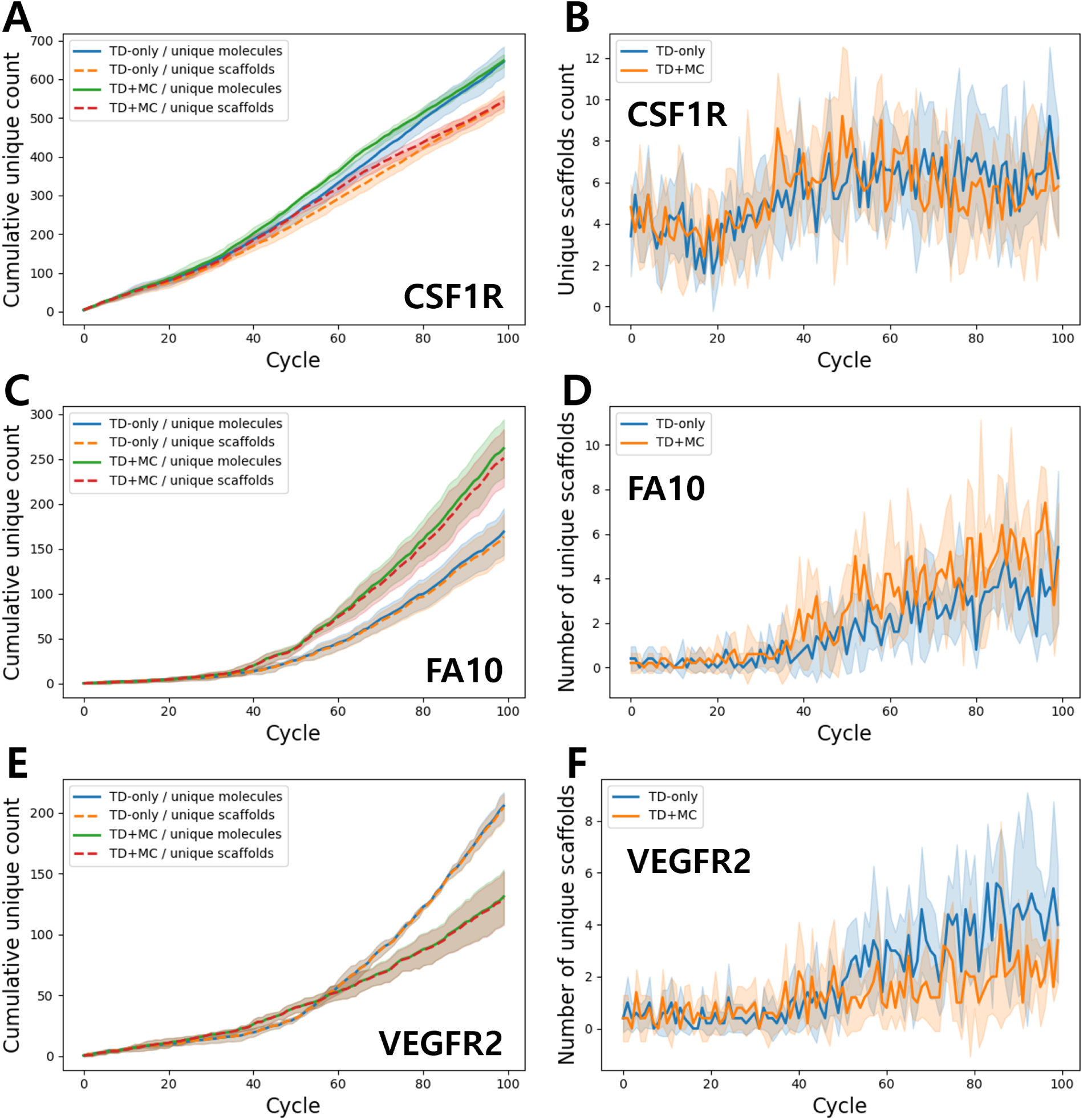
Discovery of molecules and scaffolds passing the docking score cutoff across learning cycles. Only molecules with docking scores lower than the predefined cutoff values were included: −8 kcal/mol for CSF1R, −9 kcal/mol for FA10, and −11 kcal/mol for VEGFR2. Panels A, C, and E show the cumulative numbers of unique molecules and unique scaffolds discovered across cycles for CSF1R, FA10, and VEGFR2, respectively. Panels B, D, and F show the number of unique scaffolds discovered in each cycle under the same cutoff criteria for CSF1R, FA10, and VEGFR2, respectively. Solid lines represent the mean of five independent runs, and shaded regions indicate the standard deviation.

For all targets, the cumulative numbers of cutoff-passing unique molecules and unique scaffolds increased across cycles (Figure 5A,C,E). The cycle-wise number of cutoff-passing unique scaffolds also tended to increase as learning progressed (Figure 5B,D,F). This trend was more pronounced for FA10 and VEGFR2, whereas the increase was more modest for CSF1R.

This behavior contrasts with the gradual slowdown in the overall discovery of unique molecules and scaffolds shown in Figure 4B,D,F. Although the generation of overall new molecules became less frequent in later cycles, FragDockRL continued to increase the discovery of molecules and scaffolds satisfying the docking score cutoff. This suggests that Q-guided action selection increased the enrichment of cutoff-passing candidates while the overall exploration became narrower.

We further examined the effect of multi-product BB addition filtering using the BB addition cases accumulated over FragDockRL learning cycles. As described in Methods, BB addition cases were excluded when a reaction between the current molecule and a candidate BB could produce multiple structurally distinct products, because such cases would make the resulting state ambiguous. To assess the scale of this filtering during FragDockRL searches, we counted the numbers of BB addition cases yielding a single product or multiple products during the runs.

As shown in Table 2, multi-product cases accounted for a substantial fraction of the BB addition cases observed during FragDockRL searches. The fraction was approximately 20% or higher for all tested targets and variants, and was highest for CSF1R, reaching about 50%. Thus, the multi-product filter restricted a non-negligible portion of the accessible BB addition cases. This restriction may affect the accessible molecular diversity by limiting the search to BB addition cases with unambiguous product outcomes.

**Table 2:** Number of BB addition cases yielding a single product or multiple products during FragDockRL searches. A multi-product case refers to a BB addition that can generate multiple structurally distinct products due to multiple possible growth-vector assignments. Values are reported as mean ± standard deviation over five independent runs.

| Target | Variant | Single-product cases | Multi-product cases | Multi-product fraction (%) |
| --- | --- | --- | --- | --- |
| CSF1R | TD-only | 24527.4 ± 1422.6 | 23901.0 ± 197.8 | 49.3 |
| CSF1R | TD+MC | 20523.0 ± 627.4 | 24095.2 ± 275.9 | 54.0 |
| FA10 | TD-only | 37627.2 ± 2473.8 | 15383.6 ± 1201.6 | 29.0 |
| FA10 | TD+MC | 33814.6 ± 1524.2 | 9221.0 ± 1200.5 | 21.4 |
| VEGFR2 | TD-only | 38250.4 ± 1269.9 | 9445.0 ± 879.6 | 19.8 |
| VEGFR2 | TD+MC | 32125.4 ± 2025.9 | 8028.2 ± 900.5 | 20.0 |

### Benchmark Comparison

We next compared FragDockRL with baseline generation methods implemented within the same FragDock framework: One-Step Reaction, Random Search, BS, and MCTS. For Frag-DockRL, both TD-only and TD+MC variants were evaluated. This benchmark was intended to assess not only overall performance, but also how different strategies behave across targets with different reaction-space and docking-score characteristics. The One-Step Reaction baseline provides exhaustive enumeration of the single-step product space, whereas Random Search, BS, MCTS, and FragDockRL explore multi-step virtual synthesis trajectories under a limited molecule-generation budget. In the following discussion, we focus primarily on the TD+MC variant because it represents the intended learning setting of FragDockRL, while the TD-only variant is used as an additional comparison to examine how the training objective affects search behavior.

For each target, we compared the total number of generated molecules, the numbers of unique molecules and unique scaffolds, and the numbers of unique molecules and unique scaffolds passing the target-specific docking score cutoff. The number of unique molecules passing the docking score cutoff was used as the primary performance metric, because it directly reflects the number of candidate molecules available for subsequent selection. The same cutoff values defined in the previous subsection were used for this comparison. The results are summarized in Tables 3–5.

**Table 3:** Benchmark comparison for CSF1R. Docking scores are reported in kcal/mol, and lower values indicate more favorable docking results. The docking score cutoff was set to −8 kcal/mol. Values are reported as mean ± standard deviation over five independent runs, except for the deterministic One-Step Reaction baseline.

| Method | Generated molecules | Unique molecules | Unique scaffolds | Unique molecules score < cutoff | Unique scaffolds score < cutoff |
| --- | --- | --- | --- | --- | --- |
| One-Step Reaction | 42735 | 41380 | 12577 | 1663 | 1122 |
| Random Search | 20000 | $16011.6 \pm 33.44$ | $10660.6 \pm 35.62$ | $389.0 \pm 13.81$ | $330.8 \pm 10.78$ |
| Beam Search | 19998 | $19908.8 \pm 12.54$ | $10595.0 \pm 273.99$ | $407.4 \pm 149.44$ | $330.4 \pm 115.58$ |
| MCTS | 20000 | $17033.0 \pm 45.45$ | $11056.0 \pm 60.70$ | $388.2 \pm 15.48$ | $326.0 \pm 13.78$ |
| FragDockRL (TD-only) | 20000 | $15911.6 \pm 396.22$ | $9803.8 \pm 441.97$ | $645.0 \pm 34.53$ | $543.2 \pm 24.78$ |
| FragDockRL (TD+MC) | 20000 | $14774.2 \pm 243.45$ | $8238.4 \pm 196.74$ | $647.6 \pm 14.19$ | $542.2 \pm 12.98$ |

**Table 4:**
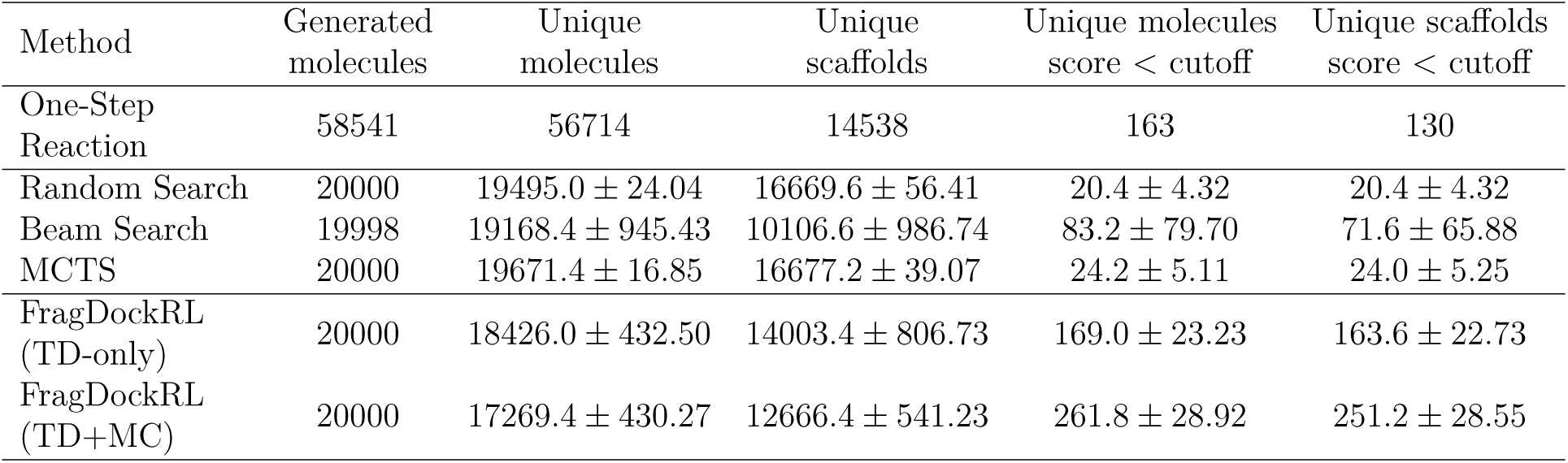
Benchmark comparison for FA10. Docking scores are reported in kcal/mol, and lower values indicate more favorable docking results. The docking score cutoff was set to −9 kcal/mol. Values are reported as mean ± standard deviation over five independent runs, except for the deterministic One-Step Reaction baseline.

| Method | Generated molecules | Unique molecules | Unique scaffolds | Unique molecules score < cutoff | Unique scaffolds score < cutoff |
| --- | --- | --- | --- | --- | --- |
| One-Step Reaction | 58541 | 56714 | 14538 | 163 | 130 |
| Random Search | 20000 | 19495.0 ± 24.04 | 16669.6 ± 56.41 | 20.4 ± 4.32 | 20.4 ± 4.32 |
| Beam Search | 19998 | 19168.4 ± 945.43 | 10106.6 ± 986.74 | 83.2 ± 79.70 | 71.6 ± 65.88 |
| MCTS | 20000 | 19671.4 ± 16.85 | 16677.2 ± 39.07 | 24.2 ± 5.11 | 24.0 ± 5.25 |
| FragDockRL (TD-only) | 20000 | 18426.0 ± 432.50 | 14003.4 ± 806.73 | 169.0 ± 23.23 | 163.6 ± 22.73 |
| FragDockRL (TD+MC) | 20000 | 17269.4 ± 430.27 | 12666.4 ± 541.23 | 261.8 ± 28.92 | 251.2 ± 28.55 |

**Table 5:** Benchmark comparison for VEGFR2. Docking scores are reported in kcal/mol, and lower values indicate more favorable docking results. The docking score cutoff was set to −11 kcal/mol. Values are reported as mean ± standard deviation over five independent runs, except for the deterministic One-Step Reaction baseline.

| Method | Generated molecules | Unique molecules | Unique scaffolds | Unique molecules score < cutoff | Unique scaffolds score < cutoff |
| --- | --- | --- | --- | --- | --- |
| One-Step Reaction | 41285 | 39616 | 10373 | 5 | 3 |
| Random Search | 20000 | 19511.0 ± 15.94 | 15005.8 ± 65.08 | 41.2 ± 5.88 | 41.2 ± 5.88 |
| Beam Search | 19998 | 19197.8 ± 364.99 | 9711.0 ± 282.99 | 1083.8 ± 763.73 | 680.6 ± 454.69 |
| MCTS | 20000 | 19870.0 ± 13.70 | 15326.4 ± 52.01 | 40.8 ± 7.93 | 40.8 ± 7.93 |
| FragDockRL (TD-only) | 20000 | 16170.2 ± 797.16 | 11939.8 ± 594.16 | 205.8 ± 9.95 | 204.4 ± 9.89 |
| FragDockRL (TD+MC) | 20000 | 18263.4 ± 256.34 | 13980.4 ± 359.82 | 131.2 ± 19.97 | 129.8 ± 19.65 |

Most search-based methods were evaluated using 20,000 generated molecules for each run. In contrast, the One-Step Reaction baseline deterministically enumerates all available single-step products from the initial state, so the number of generated molecules differs from the other methods and varies across targets. To further interpret the search behavior of each method, we analyzed the reaction-step distribution of unique molecules passing the docking score cutoff (Tables S1–S3).

Across the three targets, FragDockRL often generated fewer total unique molecules than Random Search or MCTS. This lower overall uniqueness is consistent with the focused nature of the learned policy: FragDockRL tends to prioritize BBs and molecular growth actions predicted to be valuable by the Q-network, while duplicate products are not explicitly prohibited during generation. Thus, a lower total unique count does not necessarily indicate poorer search performance, but can reflect a trade-off between broad structural coverage and enrichment of productive candidates.

For CSF1R, the One-Step Reaction baseline yielded the largest number of unique molecules passing the docking score cutoff (Table 3). Although this baseline generated more molecules than the search-based methods, it also showed the highest ratio of cutoff-passing unique molecules to generated molecules. This indicates that its strong performance was not simply due to a larger enumeration size, but reflected the high productivity of the single-step product space for this target. To further understand this result, we analyzed the reaction-step distribution of cutoff-passing unique molecules (Table S1). For all methods, cutoff-passing molecules were most frequently obtained from single-step products, suggesting that a large fraction of the accessible low-scoring molecules for CSF1R was already present in the single-step product space. Therefore, the strong performance of the One-Step Reaction baseline should not be interpreted as a general superiority of shallow enumeration, but rather as a target-dependent property of the accessible reaction space. In this case, exhaustive enumeration of single-step products was sufficient to identify a large number of favorable candidates, while also retaining scaffold-level diversity, as reflected by the number of cutoff-passing unique scaffolds.

Nevertheless, among the search-based methods, FragDockRL produced the largest number of cutoff-passing unique molecules for CSF1R. Notably, this advantage was also observed when only single-step products were compared (Table S1). This result suggests that Frag-DockRL can prioritize productive BB selections more effectively than non-learning-based search methods. In non-learning-based search methods, the usefulness of a BB is primarily assessed through the docking results of explicitly generated molecules, whereas FragDockRL learns from previously evaluated molecular structures and can estimate the potential value of candidate BBs and molecular growth actions before explicitly evaluating the corresponding docked products. Thus, even in a target where single-step products are already highly favorable, the learned policy improved the efficiency of selecting productive molecular growth actions under a limited search budget.

FA10 showed a different behavior and provided the clearest example in this benchmark where learning-guided exploration was beneficial (Table 4). Although the One-Step Reaction baseline produced a non-negligible number of cutoff-passing molecules, this result was obtained by exhaustive enumeration of a much larger single-step product set, whereas the search-based methods were limited to 20,000 generated molecules. Indeed, the ratio of cutoff-passing unique molecules to generated molecules was higher for FragDockRL (TD+MC) than for the One-Step Reaction baseline. Under this limited search setting, FragDockRL (TD+MC) achieved the best performance in terms of the number of unique molecules passing the docking score cutoff.

The reaction-step distribution further supports this interpretation (Table S2). For FA10, cutoff-passing molecules were mainly obtained from two-step products, indicating that productive candidates were not confined to the shallow single-step product space. FragDockRL (TD+MC) generated the largest number of cutoff-passing molecules in both the two-step and three-step product categories, suggesting that the learned policy helped guide the search toward productive multi-step synthesis trajectories. Among the three benchmark targets, FA10 most clearly demonstrates the advantage of FragDockRL, because successful candidate discovery benefited from exploration beyond the single-step product space and from selective prioritization within a limited search budget.

VEGFR2 showed a distinct behavior from the other targets. In this case, BS produced the largest number of unique molecules passing the docking score cutoff, producing a substantially larger number than the other methods (Table 5). The reaction-step distribution provides additional insight into this result (Table S3). BS generated large numbers of cutoff-passing molecules from both two-step and three-step products, suggesting that early high-scoring branches frequently led to productive molecules after further extension. In general, this behavior is not guaranteed in multi-step molecular growth, because adding another BB can disrupt the binding pose or introduce steric clashes and other unfavorable interactions within the binding pocket. Therefore, the strong performance of BS for VEGFR2 suggests that this benchmark case was particularly well matched to a greedy expansion strategy. However, the different trends observed for CSF1R and FA10 indicate that this behavior may not be readily generalizable across targets.

Although FragDockRL did not outperform BS for VEGFR2, it still generated substantially more cutoff-passing molecules than Random Search and MCTS. This suggests that the learned policy improved the search compared with these non-learning-based methods. Interestingly, the TD-only variant outperformed the TD+MC variant for VEGFR2, in contrast to FA10 where TD+MC showed the best performance. This suggests that the relative benefit of including MC returns may depend on the target-specific search characteristics.

Overall, the benchmark results show that the most effective search strategy differed across targets, as discussed above. Because molecule-level uniqueness alone does not fully capture structural diversity among selected candidates, we also reported the number of cutoff-passing unique scaffolds as a complementary measure. This metric was used to assess whether docking-score-based filtering retained multiple scaffold families rather than selecting only closely related analogs. Across the benchmark, cutoff-passing unique scaffold counts were lower than cutoff-passing unique molecule counts, as expected, but they represented a substantial fraction of the selected molecules. For FragDockRL (TD+MC), the scaffold-to-molecule ratios among cutoff-passing candidates were 83.7%, 96.0%, and 98.9% for CSF1R, FA10, and VEGFR2, respectively.

We further compared Random Search and MCTS with FragDockRL, as these baselines were not discussed in detail in the target-specific comparisons. Random Search serves as an unguided baseline that samples candidate BBs and molecular growth actions without explicitly prioritizing predicted high-reward actions. FragDockRL uses the same virtual synthesis setting and molecule-generation budget, but biases BB selection and molecular growth through the learned Q-network. Therefore, the consistent improvement of FragDockRL over Random Search across all three targets suggests that the Q-network learned useful patterns for prioritizing productive BB selections and molecular growth actions.

MCTS did not show a clear advantage over Random Search in this benchmark. This should not be interpreted as a general limitation of MCTS itself, but rather as a consequence of the very large BB action space and the limited molecule-generation budget used here. Under this setting, MCTS may have spent much of its budget expanding new child nodes near the root rather than repeatedly revisiting and refining deeper branches, potentially making its behavior similar to broad stochastic sampling.

The computational time for each method was recorded for each target and is provided in Tables S4–S6 as practical runtime information under the benchmark setting. These run-time measurements were obtained on a desktop computer equipped with an AMD Ryzen 9 9950X3D CPU (16 cores, 32 threads), 64 GB DDR5 memory, and an NVIDIA GeForce RTX 5080 GPU.

### Representative Molecular Case Studies

To further examine the generated candidates at the molecular level, we analyzed representative molecules produced by FragDockRL. For CSF1R and FA10, representative molecules were selected from the FragDockRL (TD+MC) results, whereas for VEGFR2, they were selected from the FragDockRL (TD-only) results, which showed better performance than the TD+MC variant for this target. In each case, molecules were selected from the first run among the five independent runs. For each target, four generated molecules were selected from candidates with docked-core RMSD values below 0.5 Å relative to the reference core structure, prioritizing molecules with the most favorable docking scores. This analysis was intended to assess whether the selected molecules retained binding poses consistent with the reference ligand while introducing structural variation through different BB combinations.

Representative molecules for CSF1R, FA10, and VEGFR2 are shown in Figures 6, 7, and 8, respectively, together with their docking poses and 2D structures. Detailed information for the selected generated molecules, including the BB SMILES, reaction names, final generated molecule SMILES, docking scores, docked-core RMSDs relative to the reference core structure, and Tanimoto similarities to the reference ligands, is provided in Tables S7–S9. Tanimoto similarities were calculated using 1024-bit Morgan fingerprints with radius 2. The BBs used during FragDockRL generation and the corresponding final products are summarized in Figures S1–S3.

**Figure 6:**
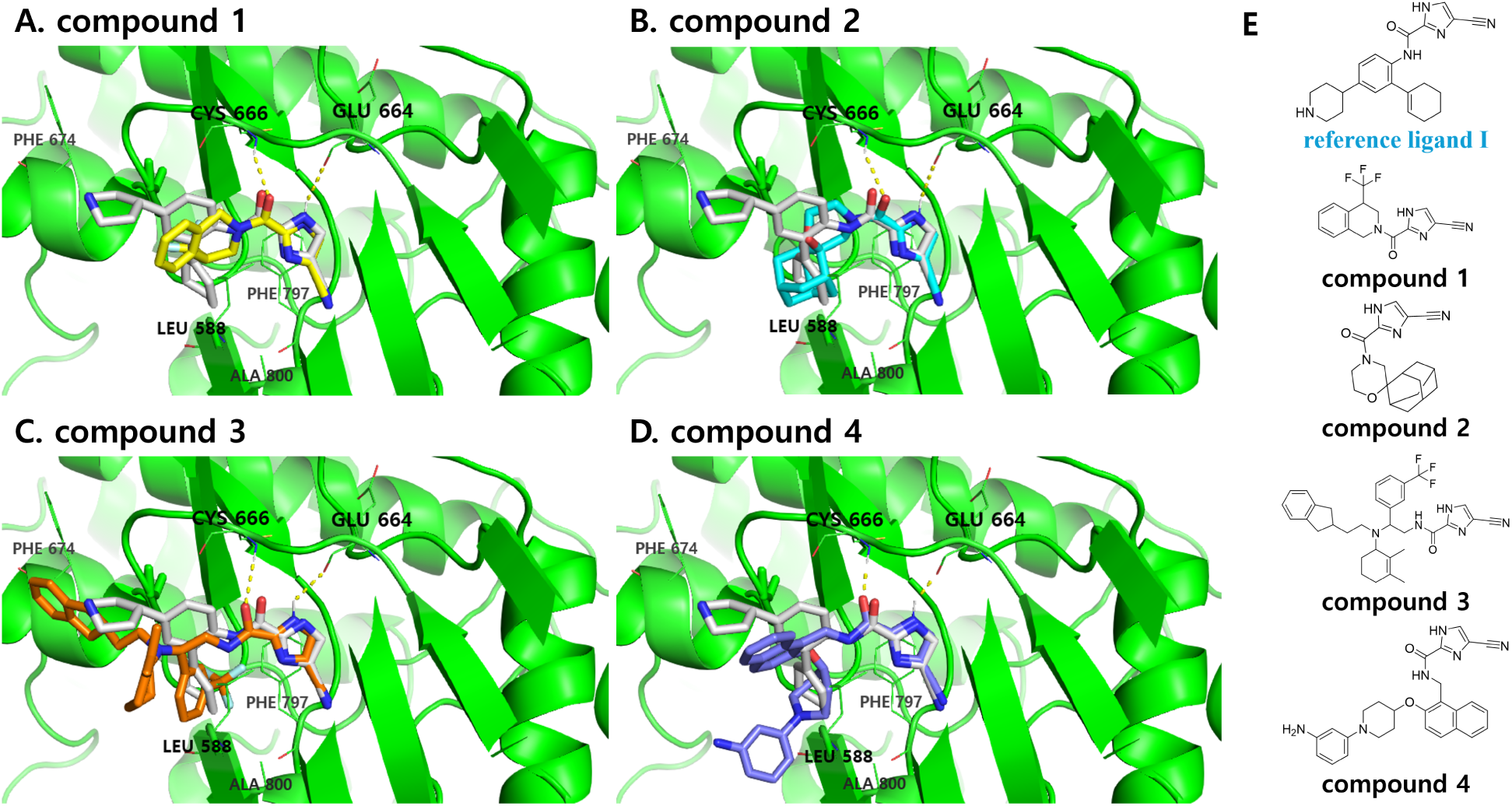
Representative generated molecules and docking poses for CSF1R. (A–D) Docking poses of compounds **1–4**, respectively, overlaid with the reference ligand **I**; (E) 2D structures of compounds **1–4** and the reference ligand **I**. Compounds **1–4** are shown in yellow, cyan, orange, and blue, respectively, and the reference ligand **I** is shown in gray. Docking score and docked-core RMSD values are as follows: compound **1**, −9.62 kcal/mol (0.38 Å); compound **2**, −9.58 kcal/mol (0.45 Å); compound **3**, −9.45 kcal/mol (0.46 Å); compound **4**, −9.34 kcal/mol (0.30 Å); and reference ligand **I**, −8.56 kcal/mol.

**Figure 7:**
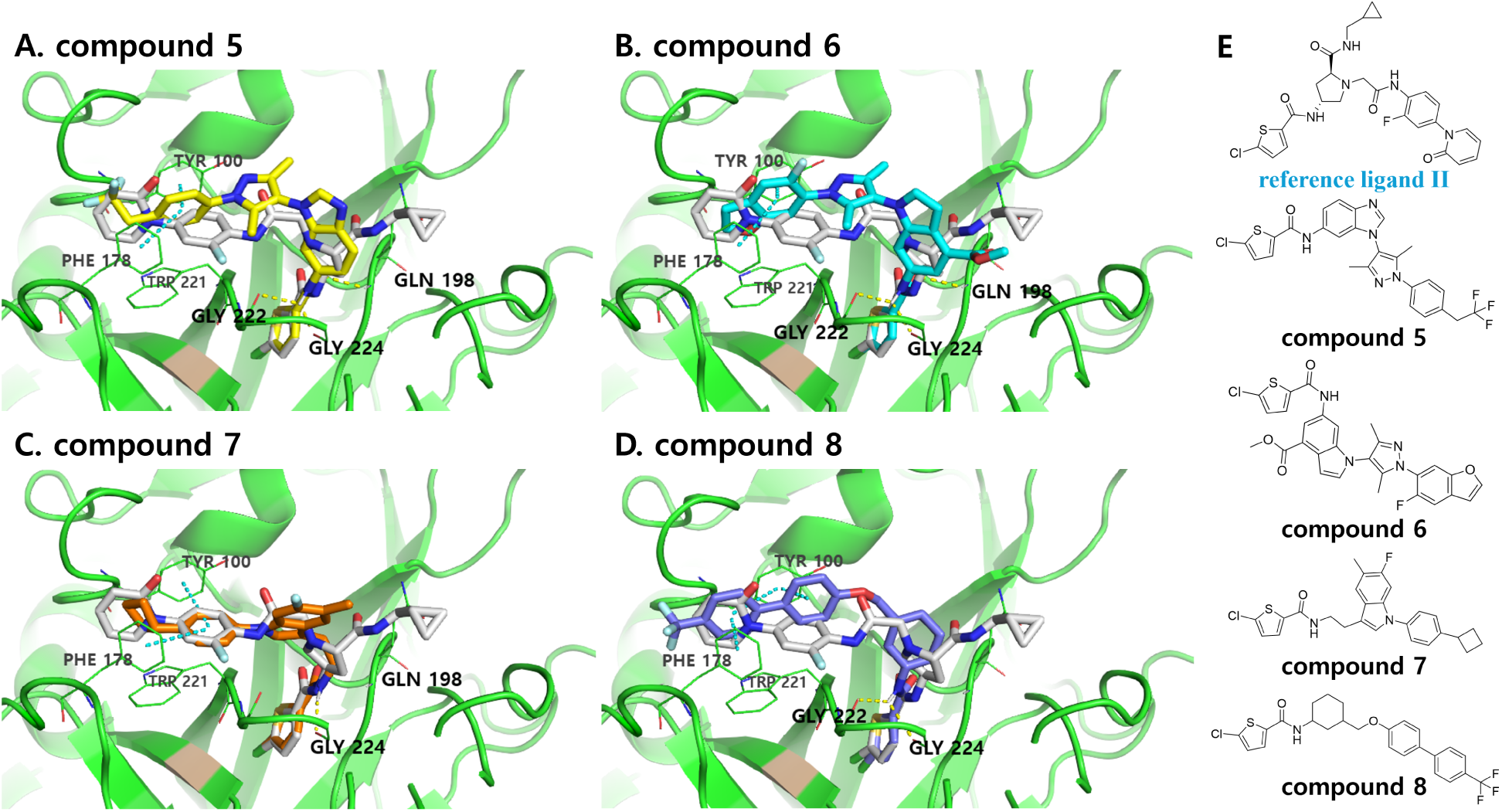
Representative generated molecules and docking poses for FA10. (A–D) Docking poses of compounds **5–8**, respectively, overlaid with the reference ligand **II**; (E) 2D structures of compounds **5–8** and the reference ligand **II**. Compounds **5–8** are shown in yellow, cyan, orange, and blue, respectively, and the reference ligand **II** is shown in gray. Docking score and docked-core RMSD values are as follows: compound **5**, −11.54 kcal/mol (0.48 Å); compound **6**, −11.22 kcal/mol (0.47 Å); compound **7**, −11.10 kcal/mol (0.46 Å); compound **8**, −11.00 kcal/mol (0.35 Å); and reference ligand **II**, −10.14 kcal/mol.

**Figure 8:**
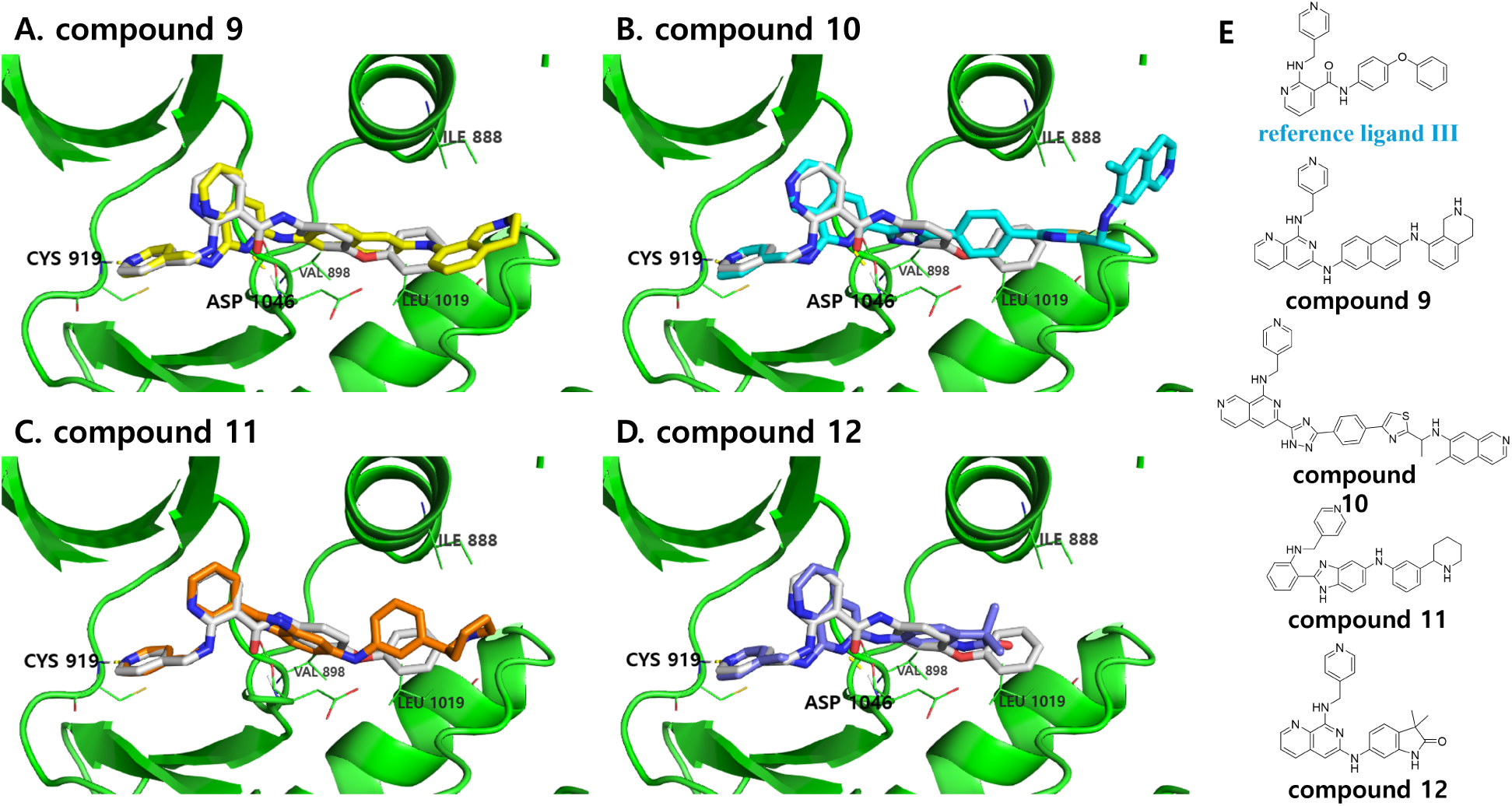
Representative generated molecules and docking poses for VEGFR2. (A–D) Docking poses of compounds **9–12**, respectively, overlaid with the reference ligand **III**; (E) 2D structures of compounds **9–12** and the reference ligand **III**. Compounds **9–12** are shown in yellow, cyan, orange, and blue, respectively, and the reference ligand **III** is shown in gray. Docking score and docked-core RMSD values are as follows: compound **9**, −13.24 kcal/mol (0.20 Å); compound **10**, −12.98 kcal/mol (0.36 Å); compound **11**, −12.67 kcal/mol (0.26 Å); compound **12**, −12.64 kcal/mol (0.30 Å); and reference ligand **III**, −11.42 kcal/mol.

The X-ray co-crystal structure of reference ligand **I** in complex with CSF1R (PDB: 3KRJ) and the docked poses of compounds **1–4** indicate that the 2-carbonyl-1H-imidazole-5-carbonitrile moiety is conserved and adopts a similar orientation within the binding site, forming consistent hydrogen bonds with Glu664 and Cys666 (Figure 6). In reference ligand **I**, the 2-cyclohexenyl group is well accommodated within a hydrophobic pocket composed of Leu588, Phe797, and Ala800. Correspondingly, the adamantane group of compound **2**, the 2,3-dimethylcyclohex-2-ene group of compound **3**, and the N-phenylpiperidine group of compound **4** occupy the same region, contributing to favorable nonpolar interactions. Despite its relatively smaller size, the 4-(trifluoromethyl)-1,2,3,4-tetrahydroisoquinoline moiety of compound **1** also effectively occupies this hydrophobic pocket. Overall, the conservation of key hydrogen-bonding interactions and hydrophobic pocket occupancy suggests that compounds **1–4** retain key recognition elements associated with CSF1R binding.

Consistent with this, compounds **1–4** exhibit improved docking scores (−9.62, −9.58, −9.45, and −9.34 kcal/mol, respectively) compared to reference ligand I (−8.56 kcal/mol), supporting their selection as representative candidates for further inspection. The Tanimoto similarities of compounds **1–4** to reference ligand I are 0.30, 0.27, 0.32, and 0.33, respectively. Together with their improved docking scores, these moderate similarity values suggest that the FragDockRL-generated compounds retain reference-like binding interactions while occupying structurally distinct chemical space.

The X-ray co-crystal structure of reference ligand **II** in complex with FA10 (PDB: 2VWM) and the docked poses of compounds **5–8** reveal that the common 2-chlorothiophenylamide moiety is conserved, occupying the deep binding pocket of FA10 and forming a hydrogen bond with Gly224 (Figure 7). The remaining portions of the molecules extend into a shallow sub-pocket, where they engage in *π*–*π* stacking interactions with aromatic residues such as Tyr100, Trp221, and Phe178. In reference ligand II, the 1-(3-fluorophenyl)-2(1H)-pyridone group is optimally positioned within the shallow pocket, forming multiple *π*–*π* stacking interactions. Similarly, the corresponding aromatic moieties of compounds **5–8**—the (2,2,2-trifluoroethyl)phenyl group of compound **5**, the 5-fluorobenzofuran-6-yl group of compound **6**, the 4-cyclobutylphenyl group of compound **7**, and the 4^′^-(trifluoromethyl)-[1,1^′^-biphenyl] group of compound **8**—occupy this region, contributing to favorable *π*–*π* and hydrophobic interactions. Notably, compound **7** adopts a binding pose highly similar to that of reference ligand **II**. Overall, the conserved hydrogen-bonding interaction and *π*–*π* stacking network suggest that compounds **5–8** retain key recognition elements associated with FA10 binding.

Consistent with these observations, compounds **5–8** exhibit improved docking scores (−11.54, −11.22, −11.10, and −11.00 kcal/mol, respectively) compared to reference ligand **II** (−10.14 kcal/mol), supporting their selection as representative candidates for further inspection. The Tanimoto similarities of compounds **5–8** to reference ligand **II** are 0.28, 0.25, 0.36, and 0.32, respectively. Together with their improved docking scores, these moderate similarity values suggest that the FragDockRL-generated compounds retain reference-like binding interactions while occupying structurally distinct chemical space.

The X-ray co-crystal structure of reference ligand **III** in complex with VEGFR2 (PDB: 2P2I) reveals that the 2-((pyridin-4-ylmethyl)amino)nicotinyl moiety extends into the binding pocket, forming hydrogen bonds with the backbone amides of Cys919 and Asp1046 (Figure 8). The 4-phenoxyphenyl group occupies a hydrophobic pocket defined by Ile888, Leu889, Val898, and Leu1019. Docking and structural comparisons indicate that reference ligand **III** and compounds **9–12** adopt similar orientations within the binding site, preserving key binding features. Compounds **9**, **10**, and **12** form hydrogen bonds with both Cys919 and Asp1046, whereas compound **11** interacts only with Cys919. In addition, the aromatic rings of compounds **9–12** occupy the same hydrophobic pocket as the 4-phenoxyphenyl group of reference ligand III, contributing to favorable nonpolar interactions. Overall, the preservation of key hydrogen-bonding interactions and hydrophobic pocket occupancy suggests that compounds **9–12** retain key recognition elements associated with VEGFR2 binding.

Consistent with these observations, compounds **9–12** exhibit improved docking scores (−13.24, −12.98, −12.67, and −12.64 kcal/mol, respectively) compared to reference ligand **III** (−11.42 kcal/mol), supporting their selection as representative candidates for further inspection. The Tanimoto similarities of compounds **9–12** to reference ligand **III** are 0.32, 0.21, 0.23, and 0.33, respectively. Together with their improved docking scores, these moderate similarity values suggest that the FragDockRL-generated compounds retain reference-like binding interactions while occupying structurally distinct chemical space.

The BBs and reaction schemes for each molecule are presented in Schemes S1–S12. As shown in these schemes, the selected compounds **1–12** are synthetically accessible from commercially available BBs in up to three steps, with an average of 2.3 steps. The synthetic routes primarily involve well-established transformations, including amination, reductive amination, Buchwald coupling, Chan–Lam coupling, Mitsunobu reactions, and triazole formation. These reactions are commonly used in medicinal chemistry and are generally considered experimentally tractable. Overall, the selected FragDockRL-generated molecules are constructed from readily accessible BBs, suggesting that their synthesis can be achieved through practical and efficient routes.

## Conclusion

In this study, we developed FragDock, a molecular design framework that combines BB-based virtual synthesis with tethered docking to explore synthetically constrained molecular spaces around a predefined core structure. Within this framework, we introduced FragDockRL as a reinforcement learning-based search method that uses docking-score-based rewards and a Q-network to prioritize productive molecular growth actions under a limited molecule-generation budget.

The training-cycle analysis showed that FragDockRL progressively enriched molecules with favorable docking scores during learning. In the benchmark comparison, FragDockRL generated more cutoff-passing unique molecules than Random Search across the three targets, supporting the role of learning-guided prioritization over unguided stochastic sampling. The benchmark also showed that the best-performing search strategy differed across targets, indicating that no single search mode was universally optimal within the tested settings. Instead, the results highlight the complementary roles of different FragDock search modes: One-Step Reaction, BS, and FragDockRL each provided advantages depending on the target-specific reaction-space characteristics. Among these cases, FragDockRL showed its clearest advantage for FA10, where FragDockRL (TD+MC) produced the largest number of cutoff-passing unique molecules among the tested methods.

Beyond benchmark-level performance, the representative molecular case studies showed that the selected compounds retained reference-like binding poses while introducing structural variation in peripheral regions. In addition, the associated reaction schemes used commercially available BBs and well-established medicinal chemistry transformations in up to three steps, supporting the synthetic plausibility of the selected compounds within the FragDock framework.

Several limitations should be noted. Docking scores were used as the primary reward and evaluation metric, but they cannot substitute for experimental binding affinity or biological activity, requiring experimental validation. In addition, the performance of FragDock and FragDockRL depends on the target protein, predefined core structure, and accessible BB/reaction space, and therefore cannot be assumed to generalize uniformly to arbitrary targets or starting cores. Finally, this benchmark focused on docking-score-based optimization. Although drug-likeness-related rewards based on MW, logP, the number of hydrogen bond acceptors (NHBA), and the number of hydrogen bond donors (NHBD) have been implemented, they were not used here, leaving ADMET- and property-aware optimization as an important direction for future development.

Beyond property-aware optimization, several directions remain for future development of FragDock. The target-dependent benchmark results suggest that different search modes within FragDock are complementary, motivating hybrid search strategies that combine learning-guided action selection with One-Step Reaction or BS. Such future versions could improve search efficiency by balancing shallow enumeration, greedy expansion, and learning-guided prioritization within the same molecular generation workflow. In addition, expanding the reaction library and introducing reaction pathways that enable BB modification would al-low FragDock to explore more diverse molecular transformations, including local structural optimization around generated candidates.

## Acknowledgement

We thank Insung Na and Sungwoo Choi of Tinaclon Inc. for their valuable assistance in the service implementation and testing of this method. S. Kang acknowledges support from the National Research Foundation of Korea (NRF) under grants NRF-2022M3E5F3080873 and RS-2024-00334098.

## Supporting Information Available

Supplementary figures, schemes, tables, and additional molecular data are provided in the Supporting Information available online.

## Data and Software Availability

The source code for the FragDock framework is available for academic and non-commercial use at https://github.com/novelism/FragDock.

Commercial use requires prior permission from Seung Hwan Hong, the corresponding author responsible for licensing.

## TOC Graphic

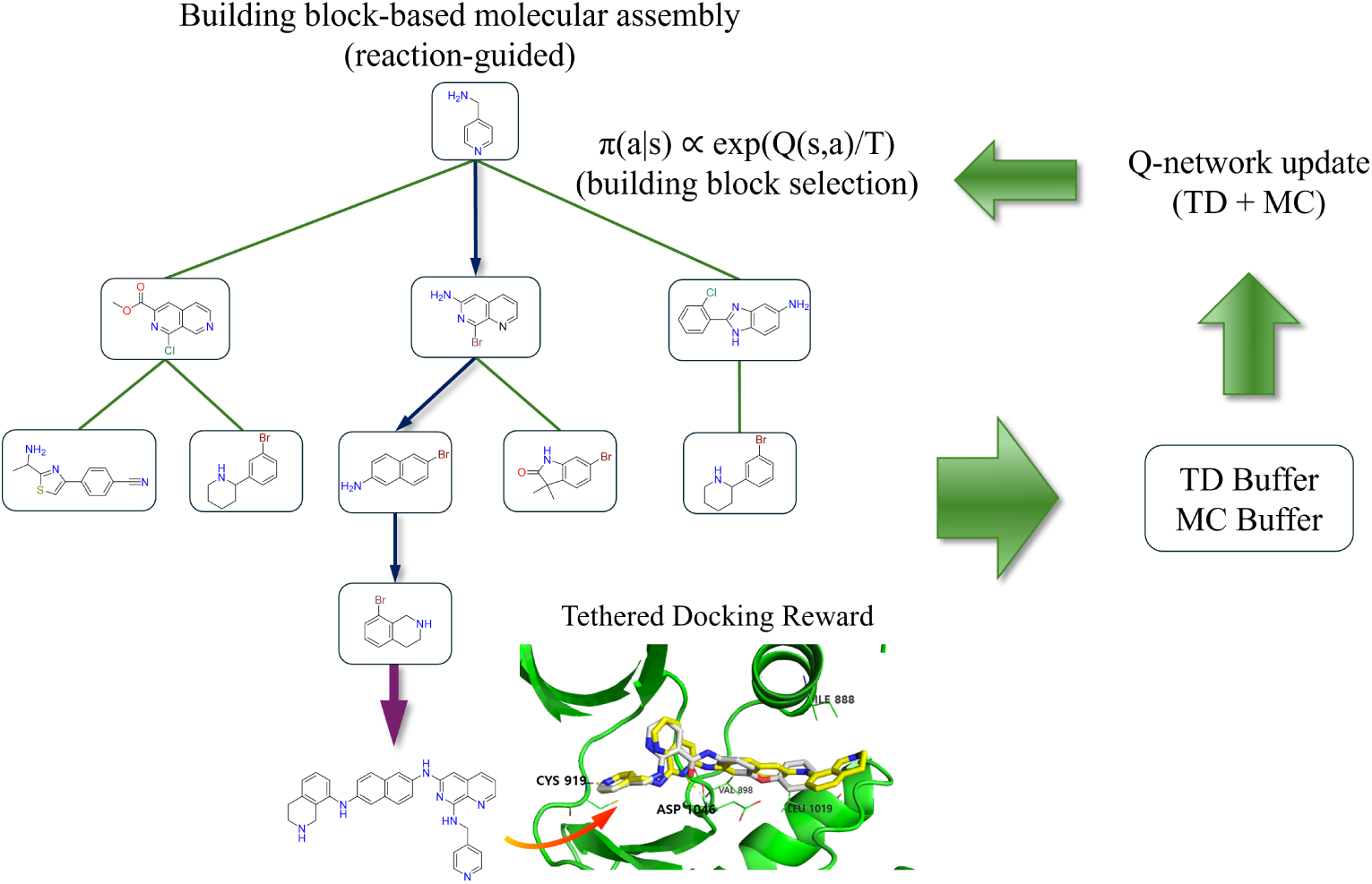

