## Supplementary Information for "FragDockRL: A Reinforcement Learning Method for Fragment-Based Ligand Design via Building Block Assembly and Tethered Docking"

#### Supplementary Tables

**Table S1.** Reaction-step distribution of unique molecules passing the docking score cutoff for CSF1R. The docking score cutoff was set to -8 kcal/mol. Here, the number of steps denotes successful virtual synthesis reactions that update the molecular state, rather than attempted molecule-generation actions. Values are reported as mean ± standard deviation over five independent runs, except for the deterministic One-Step Reaction baseline. Percentages in parentheses indicate the fraction of cutoff-passing unique molecules in each step category, calculated from the mean counts.

| Method | Unique molecules score < cutoff | 1-step product | 2-step product | 3-step product |
|---|---|---|---|---|
| One-Step Reaction | 1663 | 1663 (100.0%) | 0 (0.0%) | 0 (0.0%) |
| Random | 389.0 ± 13.8 | 363.4 ± 13.9 (93.4%) | 24.4 ± 3.9 (6.3%) | 1.2 ± 0.4 (0.3%) |
| Beam Search | 407.4 ± 149.4 | 250.4 ± 16.9 (61.5%) | 108.6 ± 154.4 (26.7%) | 48.4 ± 30.8 (11.9%) |
| MCTS | 388.2 ± 15.5 | 361.8 ± 19.8 (93.2%) | 25.8 ± 3.0 (6.6%) | 0.6 ± 1.3 (0.2%) |
| FragDockRL (TD-only) | 645.0 ± 34.5 | 488.4 ± 34.5 (75.7%) | 141.8 ± 7.3 (22.0%) | 14.8 ± 6.1 (2.3%) |
| FragDockRL (TD+MC) | 647.6 ± 14.2 | 506.8 ± 9.8 (78.3%) | 130.6 ± 11.6 (20.2%) | 10.4 ± 3.4 (1.6%) |

**Table S2.** Reaction-step distribution of unique molecules passing the docking score cutoff for FA10. Definitions and notation are the same as in Table S1.

| Method | Unique molecules score < cutoff | 1-step product | 2-step product | 3-step product |
|---|---|---|---|---|
| One-Step Reaction | 163 | 163 (100.0%) | 0 (0.0%) | 0 (0.0%) |
| Random | 20.4 ± 4.3 | 4.6 ± 1.5 (22.5%) | 14.6 ± 5.2 (71.6%) | 1.2 ± 1.3 (5.9%) |
| Beam Search | 83.2 ± 79.7 | 17.6 ± 3.4 (21.2%) | 57.4 ± 76.3 (69.0%) | 8.2 ± 14.1 (9.9%) |
| MCTS | 24.2 ± 5.1 | 5.6 ± 3.4 (23.1%) | 17.6 ± 4.8 (72.7%) | 1.0 ± 0.7 (4.1%) |
| FragDockRL | 169.0 ± 23.2 | 48.4 ± 11.5 | 101.4 ± 19.7 | 19.2 ± 2.6 |

| Method | Unique molecules score < cutoff | 1-step product | 2-step product | 3-step product |
|---|---|---|---|---|
| (TD-only) | | (28.6%) | (60.0%) | (11.4%) |
| FragDockRL (TD+MC) | 261.8 ± 28.9 | 67.6 ± 6.5 (25.8%) | 155.8 ± 36.2 (59.5%) | 38.4 ± 7.1 (14.7%) |

**Table S3.** Reaction-step distribution of unique molecules passing the docking score cutoff for VEGFR2. Definitions and notation are the same as in Table S1.

| Method | Unique molecules score < cutoff | 1-step product | 2-step product | 3-step product |
|---|---|---|---|---|
| One-Step Reaction | 5 | 5 (100.0%) | 0 (0.0%) | 0 (0.0%) |
| Random | 41.2 ± 5.9 | 0.2 ± 0.4 (0.5%) | 33.6 ± 6.8 (81.6%) | 7.4 ± 2.9 (18.0%) |
| Beam Search | 1083.8 ± 763.7 | 1.4 ± 1.5 (0.1%) | 537.8 ± 458.8 (49.6%) | 544.6 ± 500.2 (50.2%) |
| MCTS | 40.8 ± 7.9 | 0.2 ± 0.4 (0.5%) | 31.8 ± 7.5 (77.9%) | 8.8 ± 2.5 (21.6%) |
| FragDockRL (TD-only) | 205.8 ± 10.0 | 1.4 ± 1.3 (0.7%) | 114.8 ± 9.4 (55.8%) | 89.6 ± 13.7 (43.5%) |
| FragDockRL (TD+MC) | 131.2 ± 20.0 | 0.0 ± 0.0 (0.0%) | 83.8 ± 14.3 (63.9%) | 47.4 ± 12.4 (36.1%) |

**Table S4.** Computational time for each generation method in the CSF1R benchmark. Times are reported in seconds. Search time includes molecule generation, action selection, and virtual reaction application. Docking time corresponds to docking-based evaluation of generated molecules. Training time corresponds to Q-network training and is applicable only to FragDockRL. A dash indicates that the method does not include a training step. Values are reported as mean ± standard deviation over five independent runs, except for the deterministic One-Step Reaction baseline. Docking calculations were parallelized for all methods except MCTS.

| Method | Generated molecules | Search time (s) | Docking time (s) | Training time (s) |
|---|---|---|---|---|
| One-Step Reaction | 42735 | 28 | 13412 | -- |
| Random | 20000 | 130 ± 0.6 | 6997 ± 15.9 | -- |
| Beam Search | 19998 | 32 ± 7.5 | 6173 ± 584.0 | -- |
| MCTS | 20000 | 386 ± 0.2 | 145127 ± 629.9 | -- |
| FragDockRL (TD-only) | 20000 | 1110 ± 3.6 | 6510 ± 282.1 | 1635 ± 3.4 |
| FragDockRL (TD+MC) | 20000 | 1121 ± 3.3 | 5821 ± 111.8 | 1059 ± 16.4 |

**Table S5.** Computational time for each generation method in the FA10 benchmark. Definitions and notation are the same as in Table S4.

| Method | Generated molecules | Search time (s) | Docking time (s) | Training time (s) |
|---|---|---|---|---|
| One-Step Reaction | 58541 | 25 | 19357 | -- |
| Random | 20000 | 131 ± 0.7 | 7762 ± 14.4 | -- |
| Beam Search | 19998 | 16 ± 4.0 | 7150 ± 454.3 | -- |
| MCTS | 20000 | 328 ± 1.0 | 162983 ± 382.6 | -- |
| FragDockRL (TD-only) | 20000 | 876 ± 7.8 | 7792 ± 364.7 | 1098 ± 16.6 |
| FragDockRL (TD+MC) | 20000 | 819 ± 16.3 | 7110 ± 205.4 | 703 ± 14.0 |

**Table S6.** Computational time for each generation method in the VEGFR2 benchmark. Definitions and notation are the same as in Table S4.

| Method | Generated molecules | Search time (s) | Docking time (s) | Training time (s) |
|---|---|---|---|---|
| One-Step Reaction | 41285 | 19 | 8395 | -- |
| Random | 20000 | 122 ± 0.6 | 5616 ± 10.3 | -- |
| Beam Search | 19998 | 30 ± 11.6 | 4827 ± 424.9 | -- |
| MCTS | 20000 | 294 ± 0.5 | 121095 ± 549.8 | -- |
| FragDockRL (TD-only) | 20000 | 663 ± 6.6 | 6041 ± 97.5 | 940 ± 8.8 |
| FragDockRL (TD+MC) | 20000 | 597 ± 31.9 | 5334 ± 322.8 | 614 ± 13.0 |

### Supplementary Figures

#### A. Compound 1

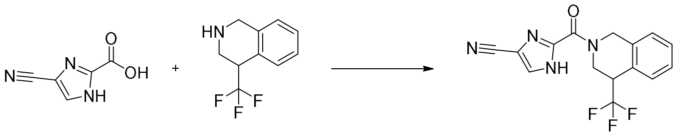

#### B. Compound 2

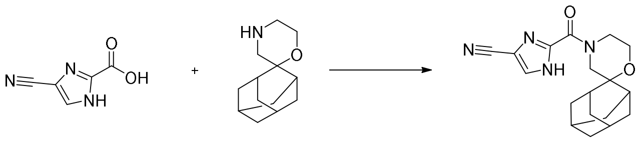

#### C. Compound 3

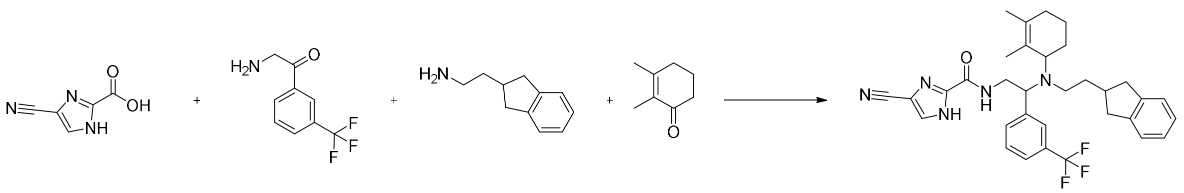

#### D. Compound 4

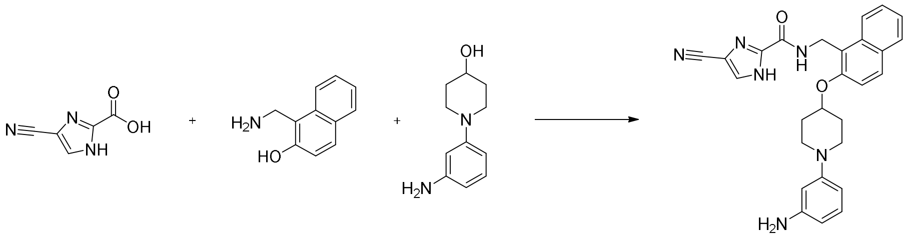

**Figure S1.** Building block composition of molecules generated by FragDockRL targeting CSF1R. Panels A-D correspond to compounds 1-4, respectively. Each panel shows the BBs selected during the FragDockRL generation process and the final generated molecule.

#### A. Compound 5

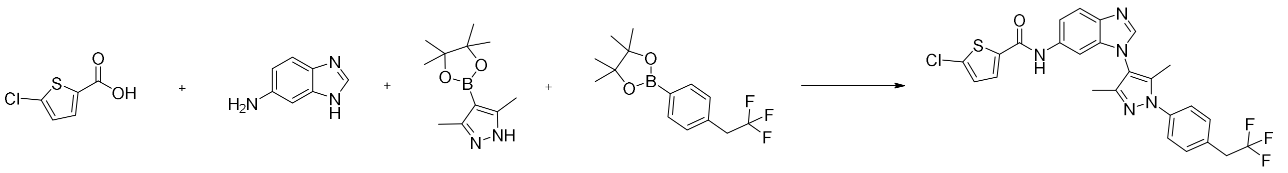

#### B. Compound 6

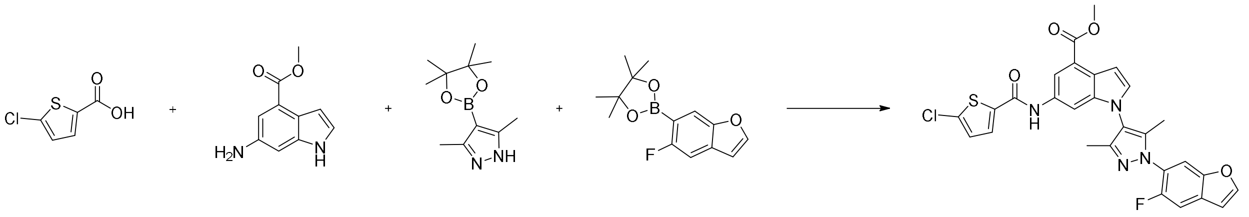

#### C. Compound 7

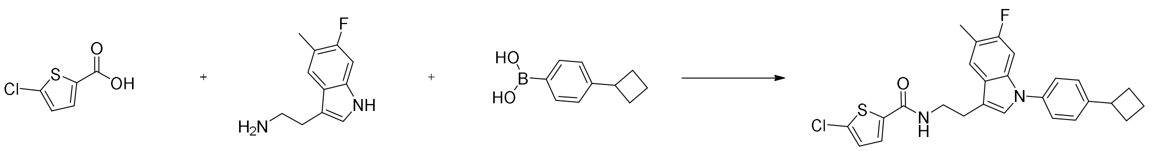

#### D. Compound 8

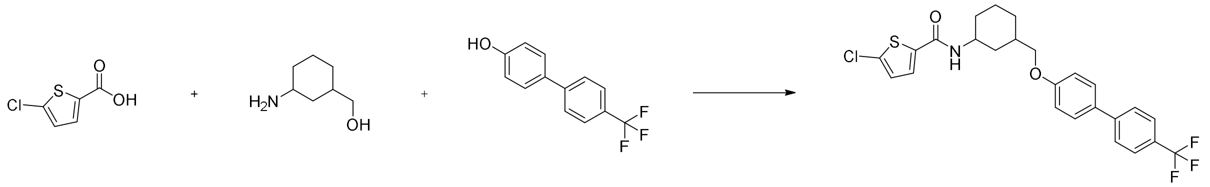

**Figure S2.** Building block composition of molecules generated by FragDockRL targeting FA10. Panels A-D correspond to compounds 1-4, respectively. Each panel shows the BBs selected during the FragDockRL generation process and the final generated molecule.

#### A. Compound 9

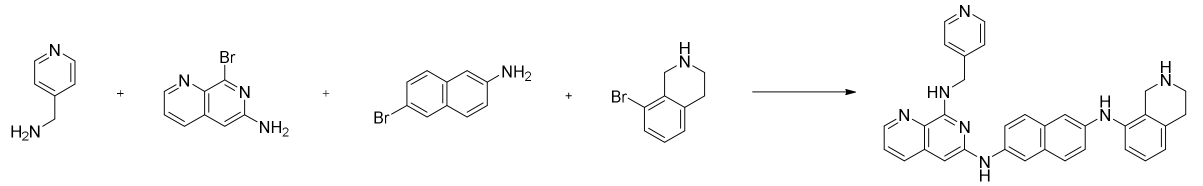


#### B. Compound 10

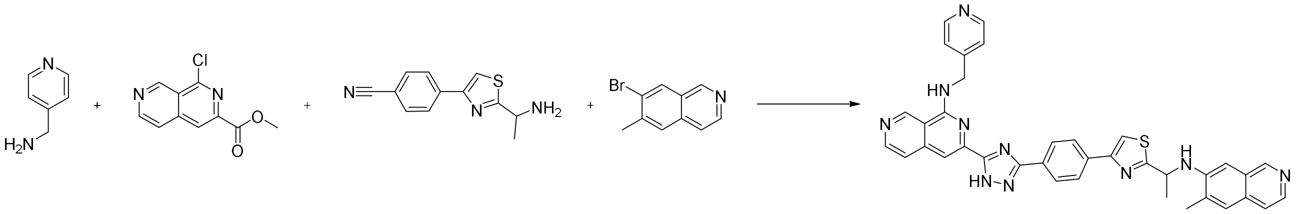


#### C. Compound 11

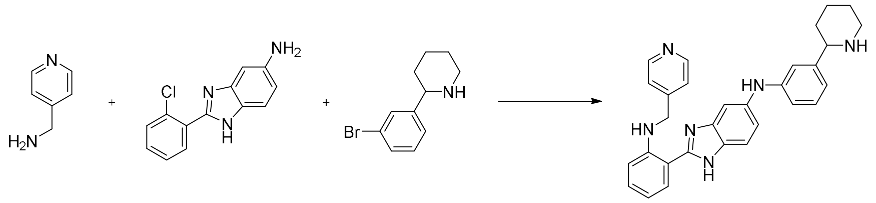


#### D. Compound 12

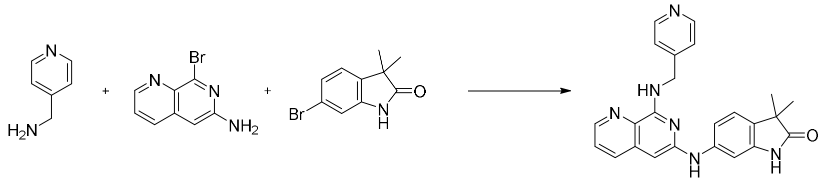


**Figure S3.** Building block composition of molecules generated by FragDockRL targeting VEGFR2. Panels A-D correspond to compounds 1-4, respectively. Each panel shows the BBs selected during the FragDockRL generation process and the final generated molecule.

### Supplementary Schemes

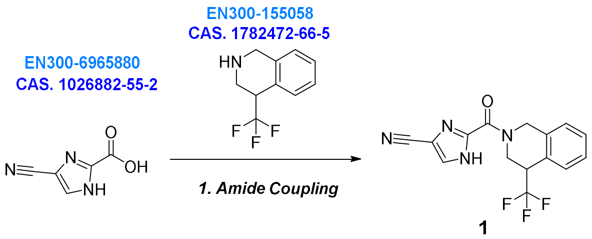


1

**Scheme S1.** Proposed synthetic route to **compound 1** from commercially available building blocks. ENAMINE building blocks are highlighted in blue by CAS number.

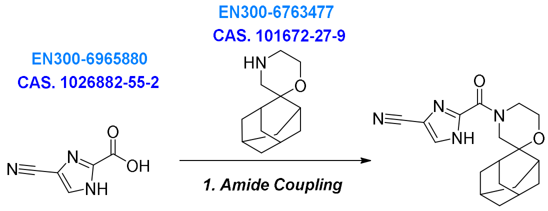


2

**Scheme S2.** Proposed synthetic route to **compound 2** from commercially available building blocks. ENAMINE building blocks are highlighted in blue by CAS number.

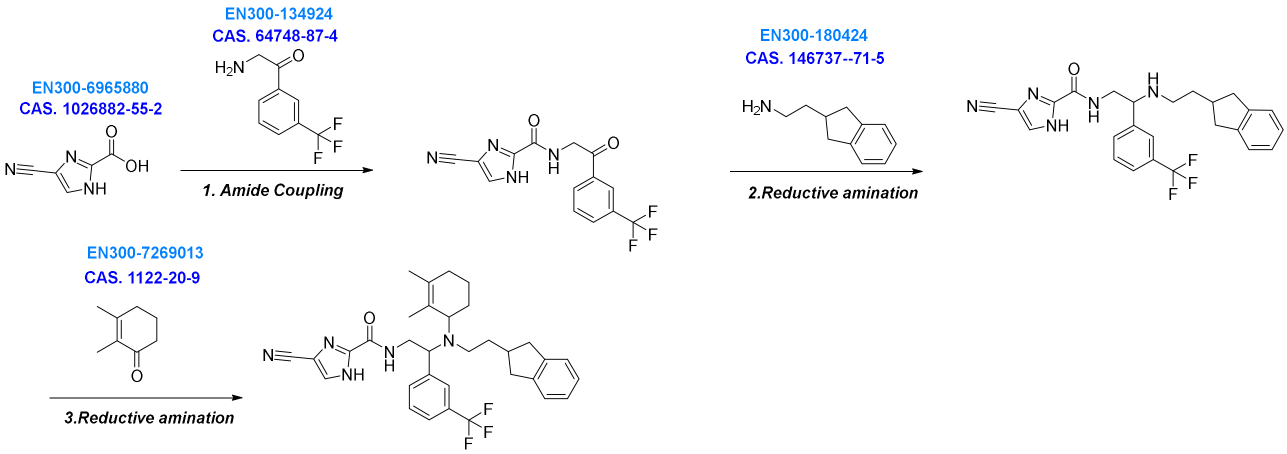


3

**Scheme S3.** Proposed synthetic route to **compound 3** from commercially available building blocks. ENAMINE building blocks are highlighted in blue by CAS number.

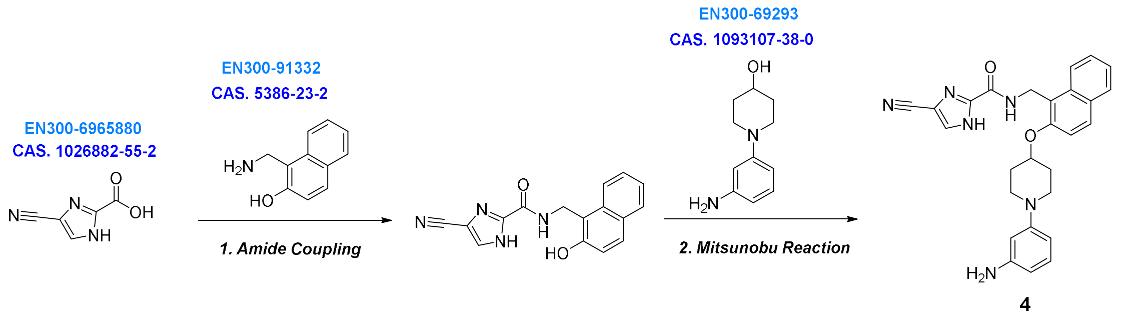


**Scheme S4.** Proposed synthetic route to **compound 4** from commercially available building blocks. ENAMINE building blocks are highlighted in blue by CAS number.

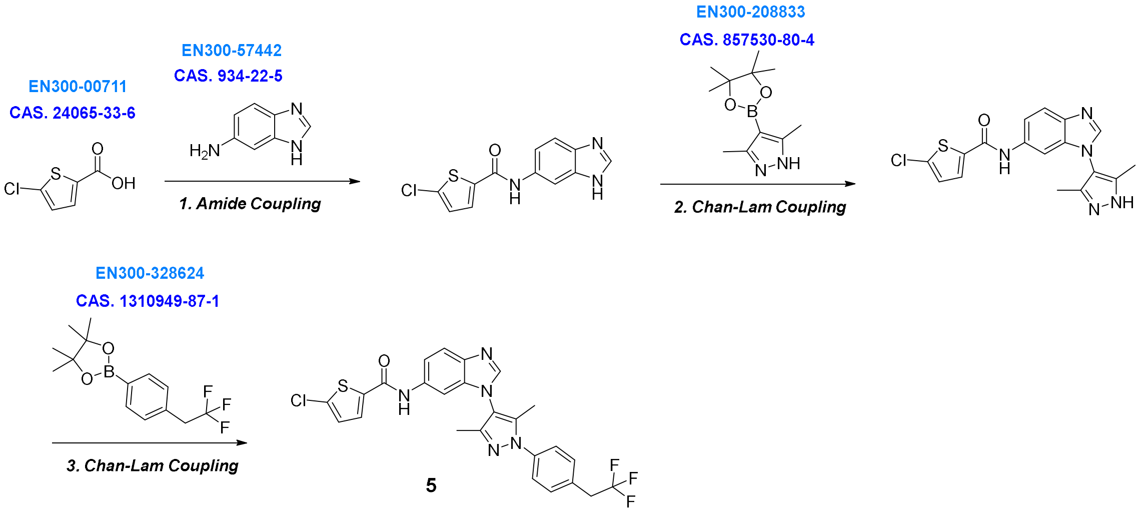


**Scheme S5.** Proposed synthetic route to **compound 5** from commercially available building blocks. ENAMINE building blocks are highlighted in blue by CAS number.

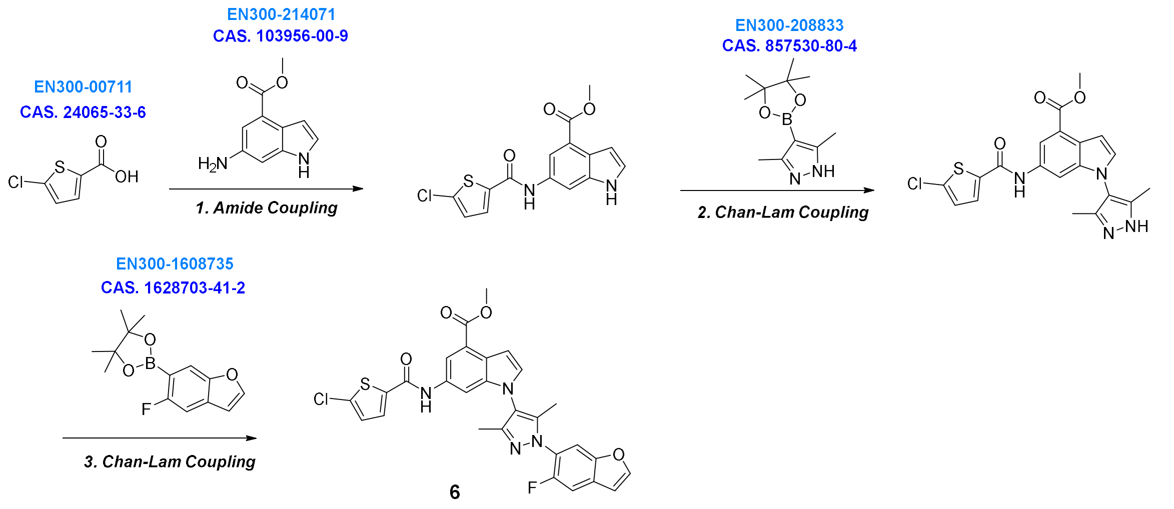


**Scheme S6.** Proposed synthetic route to **compound 6** from commercially available building blocks. ENAMINE building blocks are highlighted in blue by CAS number.

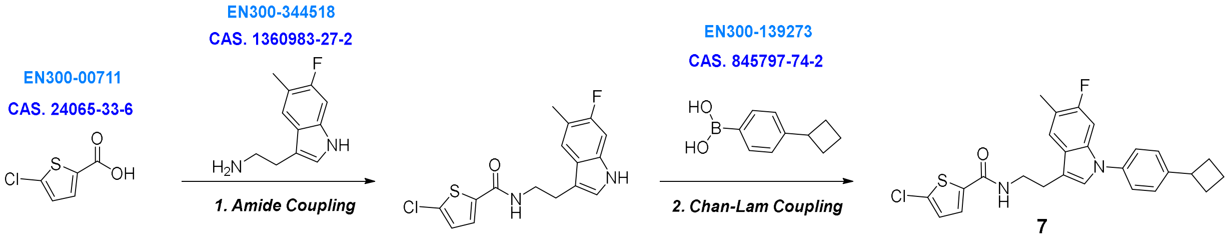


**Scheme S7.** Proposed synthetic route to **compound 7** from commercially available building blocks. ENAMINE building blocks are highlighted in blue by CAS number.

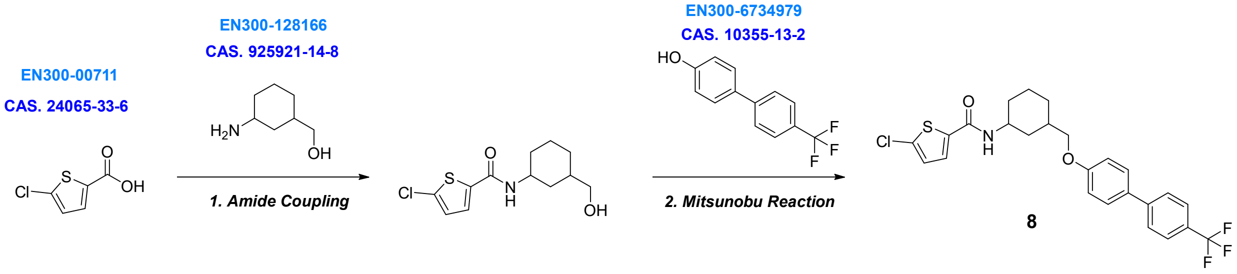


**Scheme S8.** Proposed synthetic route to **compound 8** from commercially available building blocks. ENAMINE building blocks are highlighted in blue by CAS number.

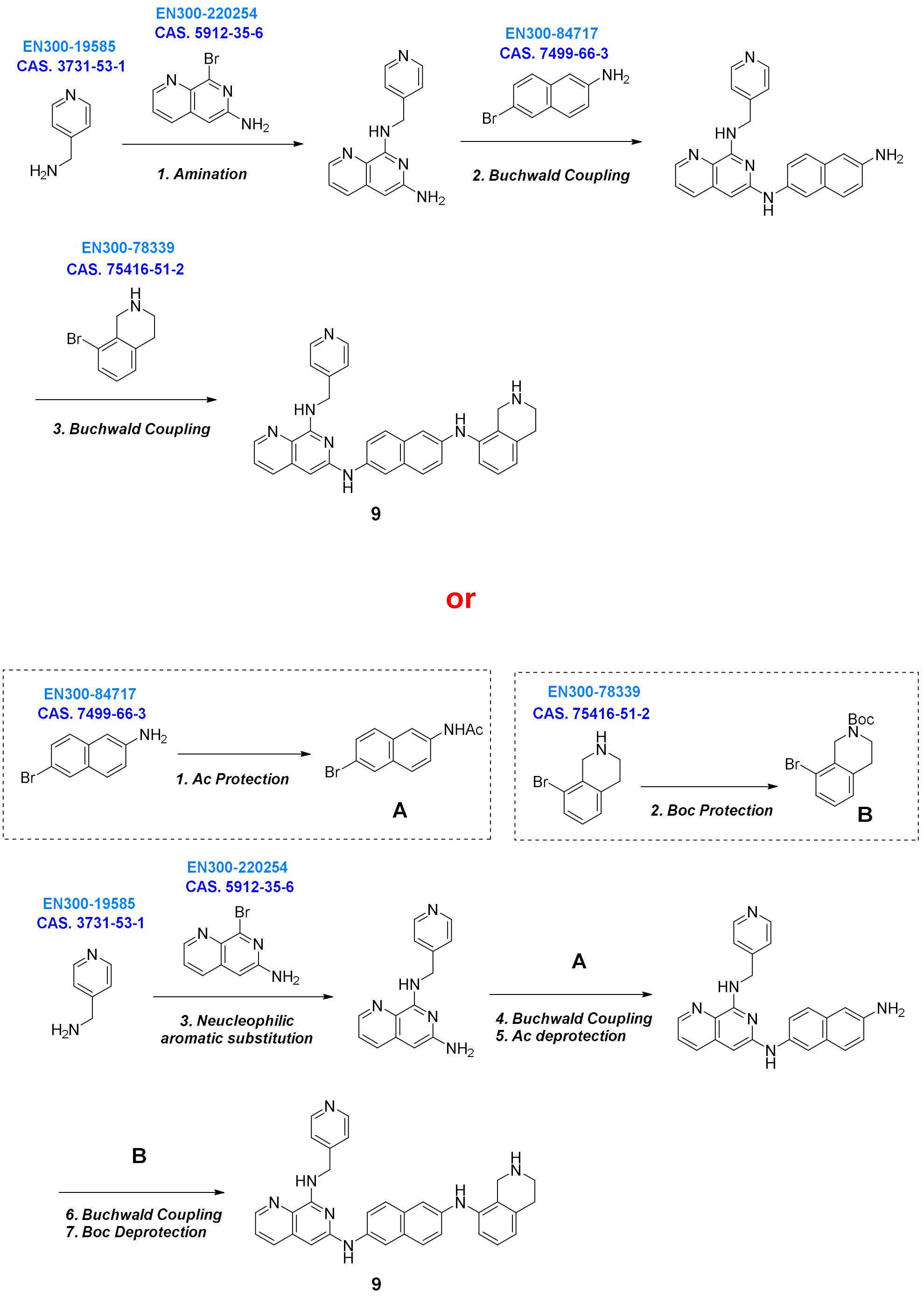


**Scheme S9.** Proposed synthetic route to **compound 9** from commercially available building blocks. ENAMINE building blocks are highlighted in blue by CAS number

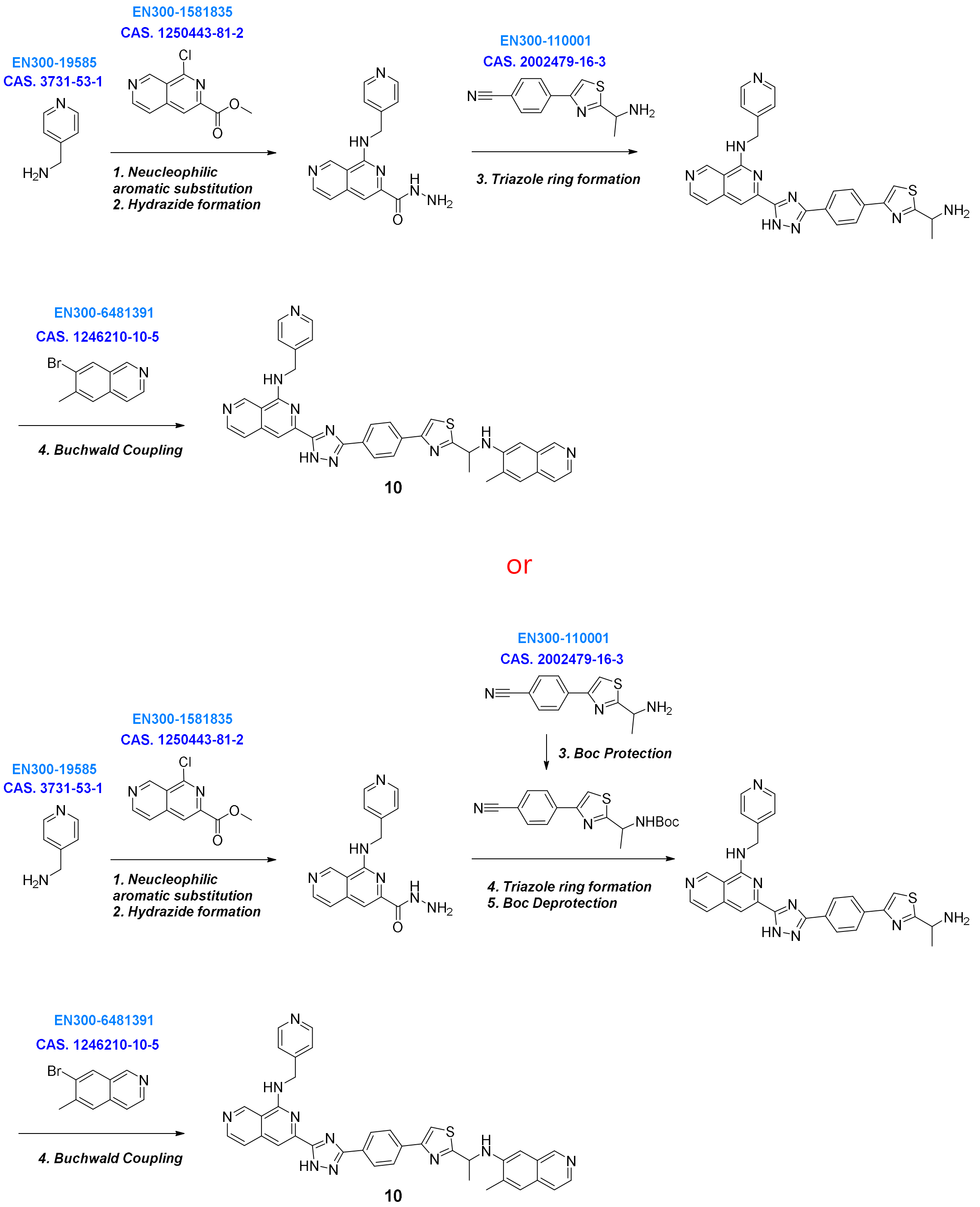


**Scheme S10.** Proposed synthetic route to **compound 10** from commercially available building blocks. ENAMINE building blocks are highlighted in blue by CAS number

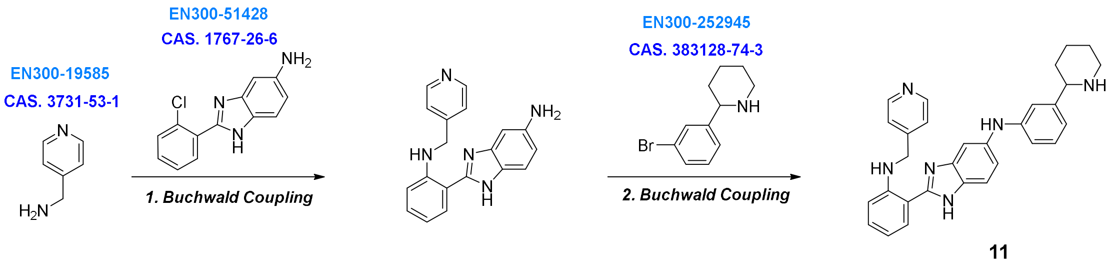


**Scheme S11.** Proposed synthetic route to **compound 11** from commercially available building blocks. ENAMINE building blocks are highlighted in blue by CAS number.

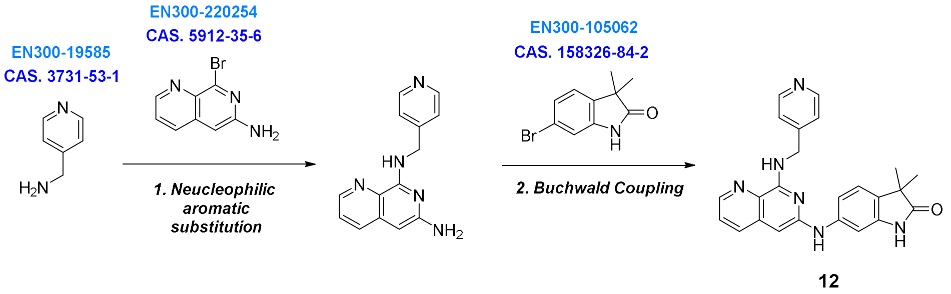


**Scheme S12.** Proposed synthetic route to **compound 12** from commercially available building blocks. ENAMINE building blocks are highlighted in blue by CAS number.
